# A non-canonical role for dynamin-1 in regulating early stages of clathrin-mediated endocytosis in non-neuronal cells

**DOI:** 10.1101/251983

**Authors:** Saipraveen Srinivasan, Christoph J. Burckhardt, Madhura Bhave, Zhiming Chen, Ping-Hung Chen, Xinxin Wang, Gaudenz Danuser, Sandra L. Schmid

## Abstract

Dynamin GTPases are best studied for their role in the terminal membrane fission process of clathrin-mediated endocytosis (CME); but, they have also been proposed to regulate earlier stages of CME. Although highly enriched in neurons, dynamin-1 (Dyn1) is, in fact, widely expressed along with dynamin-2 (Dyn2), but inactivated in non-neuronal cells via phosphorylation by GSK3β kinase. Here, we study the differential, isoform-specific functions of Dyn1 and Dyn2 as regulators of CME. Endogenously expressed Dyn1 and Dyn2 were fluorescently-tagged either separately or together in two cell lines with contrasting Dyn1 expression levels. By quantitative live cell dual and triple-channel total internal reflection fluorescence microscopy we find that Dyn2 is more efficiently recruited to clathrin-coated pits (CCPs) than Dyn1, and that Dyn2, but not Dyn1 exhibits a burst of assembly prior to CCV formation. Activation of Dyn1 by acute inhibition of GSK3β results in more rapid endocytosis of transferrin receptors, increased rates of CCP initiation and decreased CCP lifetimes, but did not significantly affect the extent of Dyn1 recruitment to CCPs. Thus, activated Dyn1 can regulate early stages of CME even when present at low, substoichiometric levels relative to Dyn2, and apparently without assembly into supramolecular collar-like structures. Under physiological conditions Dyn1 is activated downstream of EGF-receptor signaling to alter CCP dynamics. We identify sorting nexin 9 (SNX9) as a preferred binding partner to activated Dyn1 that is partially required for Dyn1-dependent effects on early stages of CCP maturation. Together, we decouple regulatory and scission functions of dynamins and report a scission-independent, isoform-specific regulatory role for Dyn1 in clathrin-mediated endocytosis.

## Introduction

Endocytosis has continued to evolve from a simple mode of ingestion and compartmentalization into a complex multicomponent process that developed a bi-directional relationship with surface signaling [1, 2]. In particular, evolutionary steps towards this complexity, which are associated with multicellularity, include the expansion to multiple isoforms of endocytic accessory proteins [3, 4], and the introduction of dynamin [4, 5].

Dynamin is the prototypical member of a family of large GTPases that catalyze membrane fission and fusion [6-8]. While encoded by single genes in *Drosophila* and *C. elegans*, further expansion of endocytic dynamins to three differentially-expressed isoforms occurred in vertebrates [9]. Dynamin-1 (Dyn1), the first identified vertebrate isoform, has been extensively studied and its mechanism of action as a fission GTPase is well understood [6, 8, 10]. The three dynamin isoforms are >70% identical in sequence, with most differences occurring in the C-terminal proline/arginine rich domain (PRD) that mediates interactions with numerous SH3 domain-containing binding partners. Dyn1 and Dyn3 appear to be functionally redundant [11]. However, Dyn2 is unable to substitute fully for Dyn1 or Dyn3 in supporting rapid synaptic vesicle recycling in neurons [12] and correspondingly, Dyn1 could not fully substitute for Dyn2 to support CME in fibroblastic cells, even when overexpressed [13]. A direct comparison of the biochemical properties of Dyn1 and Dyn2 revealed differences in their *in vitro* curvature generating abilities: Dyn1 can potently induce membrane curvature and independently catalyze vesicle release from planar membrane surfaces, whereas Dyn2 requires the synergistic activity of curvature-generating BAR domain-containing proteins [14, 15].

Less understood, and still controversial [7, 16-18] is dynamin’s suggested role in regulating early stages of clathrin-mediated endocytosis (CME) [19-22]. Based on their differential biochemical properties, it was suggested that Dyn1 might be a more effective fission GTPase, while Dyn2 might be positioned to regulate early stages of CME [14]. However, whether dynamin isoforms play distinct roles in regulating CME has not been studied.

Previously assumed to be neuron specific, recent studies have provided strong evidence that Dyn1 is indeed widely expressed, but maintained in an inactive state in non-neuronal cells through phosphorylation at S774 by the constitutively active kinase, GSK3β [23]. Acute inhibition of GSK3β in retinal pigment epithelial (ARPE) cells accelerates CME due to increased rates of clathrin-coated pit (CCP) initiation and maturation [23]. The effects of GSK3β inhibition on CME depend on Dyn1, but not Dyn2, suggesting, unexpectedly, that Dyn1 might selectively function to regulate early stages of CME in non-neuronal cells. As the GSK3 phosphorylation site, Ser774, is located within the PRD, its phosphorylation is presumed to alter interactions with dynamin’s SH3 domain-containing binding partners, as has been shown for binding partners enriched in the synapse [24, 25]. Which interactions are affected in non-neuronal cells and whether these might be dynamin isoform-specific is not known.

Immuno-electron microscopic studies using an antibody that recognizes both Dyn1 and Dyn2, have localized endogenous dynamin to both flat and deeply invaginated CCPs in A431 adenocarcinoma cells [26, 27]. Live-cell imaging has shown that, when overexpressed, both Dyn1-eGFP and Dyn2-eGFP are recruited at low levels to nascent CCPs, that their association with CCPs fluctuates and that they undergo a burst of recruitment prior to membrane scission and vesicle release [17, 22, 28-31]. Indeed, when compared directly, transiently overexpressed Dyn1‐ and Dyn2-EGFP had indistinguishable profiles for their recruitment to CCPs [30, 31]. Analysis of the recruitment of genome-edited Dyn2-eGFP to CCPs has similarly revealed a burst of recruitment at late stages of CME, as well as more transient interactions of lower numbers of Dyn2 molecules during earlier stages of CCP maturation [17, 32]. To date, direct and quantitative comparisons of the nature of Dyn1 and Dyn2 association with CCPs when they are expressed at endogenous levels do not exist. Nor is it known how activation of Dyn1 affects its association with CCPs.

Here we explore the isoform-specific behaviors of genome-edited Dyn1 and Dyn2, both at steady-state and in cells where Dyn1 is activated. We provide evidence for an early function of low levels of activtated Dyn1 in regulating CCP initiation and maturation rates and that sorting nexin 9 (SNX9) serves as an isoform-selective, and activity-dependent binding partner of Dyn1 to regulate CCP maturation. Finally, we show that Dyn1 can be activated, under physiological conditions, downstream of epidermal growth factor receptors (EGFRs) to alter CCP dynamics.

## Results

### Dynamin isoforms are differentially recruited to clathrin coated pits

Recent studies have shown that Dyn1 is widely expressed in non-neuronal cells [2]; but, like at the neuronal synapse [33], it is mostly inactive at steady state due to phosphorylation by the constitutively active kinase GSK3β. Dyn1 function and its recruitment to CCPs have been studied in non-neuronal cells, albeit under conditions of overexpression and/or without an awareness of its phospho-regulation [14, 21]. Therefore, to explore potential isoform-specific functions of Dyn1 and Dyn2, as well as the role of GSK3β in regulating Dyn1 activity, we generated genome-edited H1299 non-small cell lung cancer cells, which we previously showed partially utilize Dyn1 for CME [23]. Cells expressing endogenously-tagged Dyn2-mRuby were generated using previously validated Zinc Finger Nucleases (ZFN) [32, 34] to introduce double stranded breaks and insert the mRuby tag with complementary flanking regions by homology driven repair (**Fig 1A**). The resulting cells were single cell sorted for mRuby2 fluorescence to obtain a heterozygous clone (clone 235, designated Dyn2-mRuby^end^) expressing a single mRuby-tagged allele of Dyn2 (**Fig 1B**).

**Figure 1:**
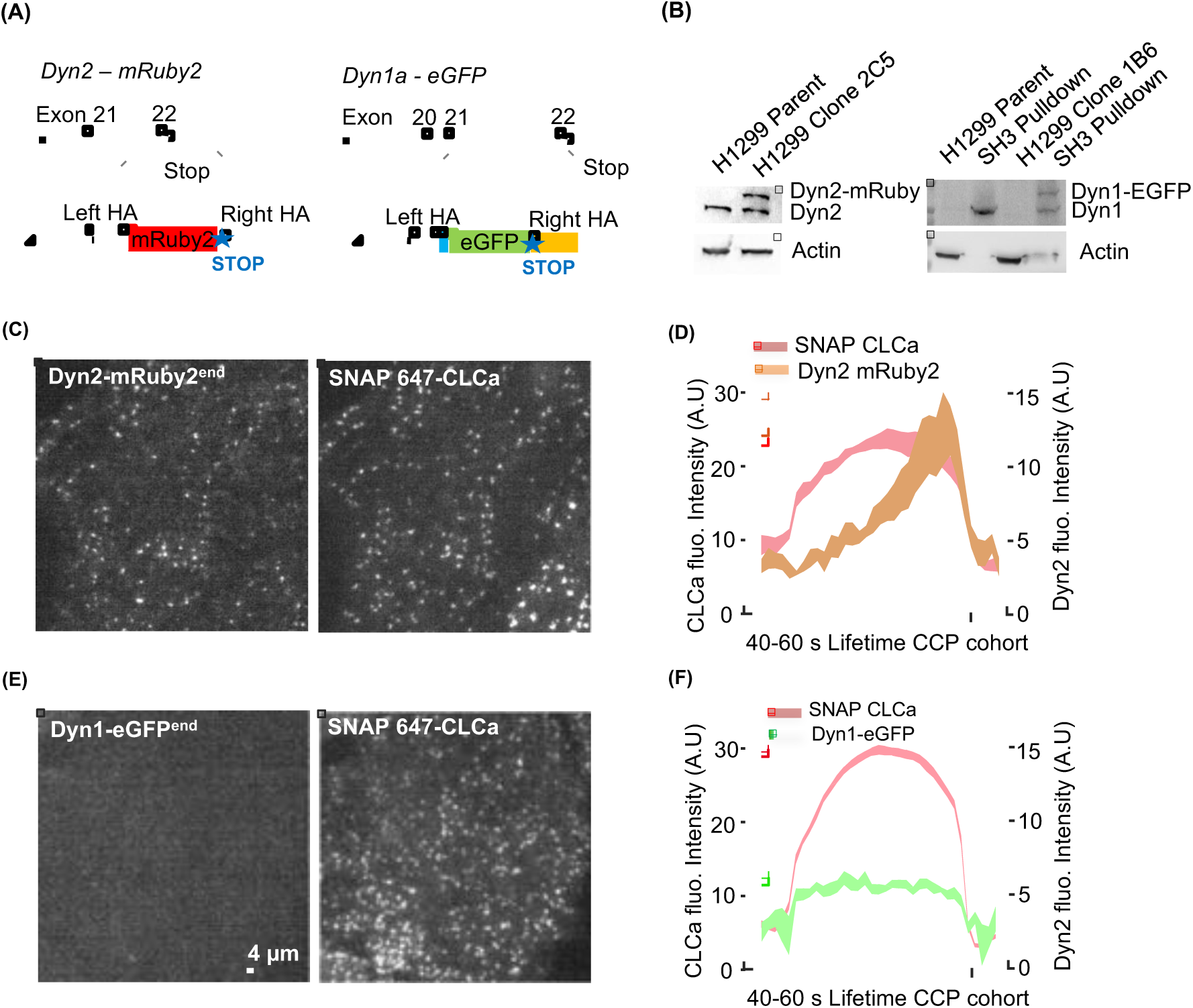
Isoform-specific differences in recruitment of dynamin to CCPs. Diagram of Zinc-finger and CRISPR-Cas9n knock-in strategies for endogenous labeling of Dyn2 and Dyn1 in H1299 cells with C-terminal mRuby2 and eGFP tags, respectively. For Dyn2, a short linker and mRuby2 (red) were placed at the stop codon in exon 22 s. For canonical Dyn1 splice isoform “a”, the 19 C-terminal amino acids (blue) were inserted in exon 21, followed by a short linker, EGFP(green) with stop codon and a polyadenylation sequence (yellow). in both constructs flanking homology arms (HA) of roughly 800bp were used to promote recombination (dashed lines). See Figure S1 for details. (B) Western blot analysis of tagged isoforms. The low levels of Dyn1 in H1299 cells could not be directly detected by western blotting, but can be detected after pulldown with GST-Amphiphysin II SH3 domains. Representative TIRF images (see Supplemental Movies S1 and Movie S2 showing membrane recruitment of endogenous Dyn2-mRuby2^end^ (C) or Dyn1a-eGFP^end^ (E) and corresponding lentiviral transduced SNAP(647)-CLCa images. (D,F) Clathrin labeled puncta were identified and thresholded to define bona fide CCPs. Shown is the averaged kinetics of recruitment of SNAP-CLCa and Dyn2-mRuby^end^ (D) or Dyn1a-eGFP^end^ (F) for all tracks with lifetimes between 40 and 60 sec (831 CCPs from 5 movies containing a total of 15 cells for Dyn2-mRuby^end^ and 13,346 Ccps from 10 movies containing a total of 29 cells for Dyn1a-eGFP^end^) is shown.

Endogenously tagging Dyn1 was complicated by the fact that the DNM1 gene encodes C-terminal splice variants derived from differential splicing of exons 21 and 22 (**Supplemental Fig S1a**), whose differential utilization could lead to partial loss of the fusion tag. Previous studies involving CRISPR/Cas9-mediated knockout and reconstitution with the Dyn1a C-terminal splice variant had confirmed that it fully reconstituted the GSK3β phospho-regulated activity of endogenous Dyn1 in H1299 cells [23], including its ability to be activated by calmodulin [35]. Therefore, using a CRISPR/Cas9n nickase strategy, we targeted the Dyn1 gene at the end of exon 21, and introduced sequences encoding for the remaining 19 amino acids of the Dyn1a isoform, followed by a seven amino acid linker [32], monomeric eGFP fusion tag with stop codon and finally the SV40 polyadenylation signal to ensure unique expression of the ‘a’ splice variant(**Fig 1A and Supplemental Fig S1**). Single cell sorting by FACS for eGFP fluorescence, followed by clonal amplification generated a heterozygous clone (clone 1B6, designated Dyn1a-eGFP^end^) expressing one eGFP-tagged allele of Dyn1a (**Fig 1B**). Note that although Dyn1 is expressed at very low levels in H1299 cells, it can be readily detected following enrichment by amphiphysin-II SH3 domain pulldown.

As a robust fiduciary marker for CCPs, clathrin light chain a (CLCa) carrying an N-terminal SNAP-fusion tag was stably introduced in parallel into both cell lines via a lentiviral vector with puromycin selection of SNAP-CLCa expressing cells. As previously reported by several groups, mild overexpression of FP-CLCa has no effect on CME as measured by transferrin endocytosis [22, 31, 32, 36], and no effect on CCP dynamics compared to AP2 or other markers [20, 29, 31]. We then performed live cell dual-channel total internal fluorescence microscopy (TIRFM) and analyzed CCP dynamics and Dyn-recruitment using the master (CLCa) – slave (Dyn) approach introduced with the cmeAnalysis software [22, 37, 38].

As expected based on previous studies using either overexpressed [28, 29, 31] or endogenously tagged Dyn2 [17, 22, 32], Dyn2-mRuby^end^ was observed, on average, to gradually accumulate and then exhibit a burst of recruitment coincident with CCV release. This can be seen in class averaged tracks of bona fide CCPs with lifetimes ranging from 40-60 sec

(**Fig 1 C, D),** and in all other CCP lifetime cohorts (**Supplemental Fig S2A,B, Supplemental movie 1**). In contrast, Dyn1a-eGFP^endo^ recruitment was barely detectable above background and no burst was evident (**Fig 1E,F, Supplemental Fig S2C,D, Supplemental movie 2**). This could reflect either isoform-specific differences, very low levels of Dyn1 expression relative to Dyn2, and/or the inactivation of Dyn1 by GSK3 phosphorylation. Thus, we further explored these possibilities.

### Inhibition of constitutively active GSK3β kinase stimulates Dyn1 to accelerate CCP initiation and maturation

We first tested whether activation of Dyn1 alters CCP dynamics and/or the recruitment of Dyn1a-eGFP^endo^ in H1299 cells. As expected based on earlier studies in ARPE cells [23], we confirmed that acute inhibition of GSK3β by incubation with the specific inhibitor, CHIR99021, leads to decreased phosphorylation of Dyn1 at Ser774 within 30 minutes (**Fig. 2A, B**) and increased rates of CME, as measured by transferrin receptor (TfnR) internalization (**Fig. 2C**). Importantly, the effects of GSK3β inhibition were dependent on Dyn1 expression, as treatment of Dyn1 knockout (Dyn1^KO^) H1299 cells [23] with CHIR99021 had no effect on CME (**Fig. 2C**).

**Figure 2:**
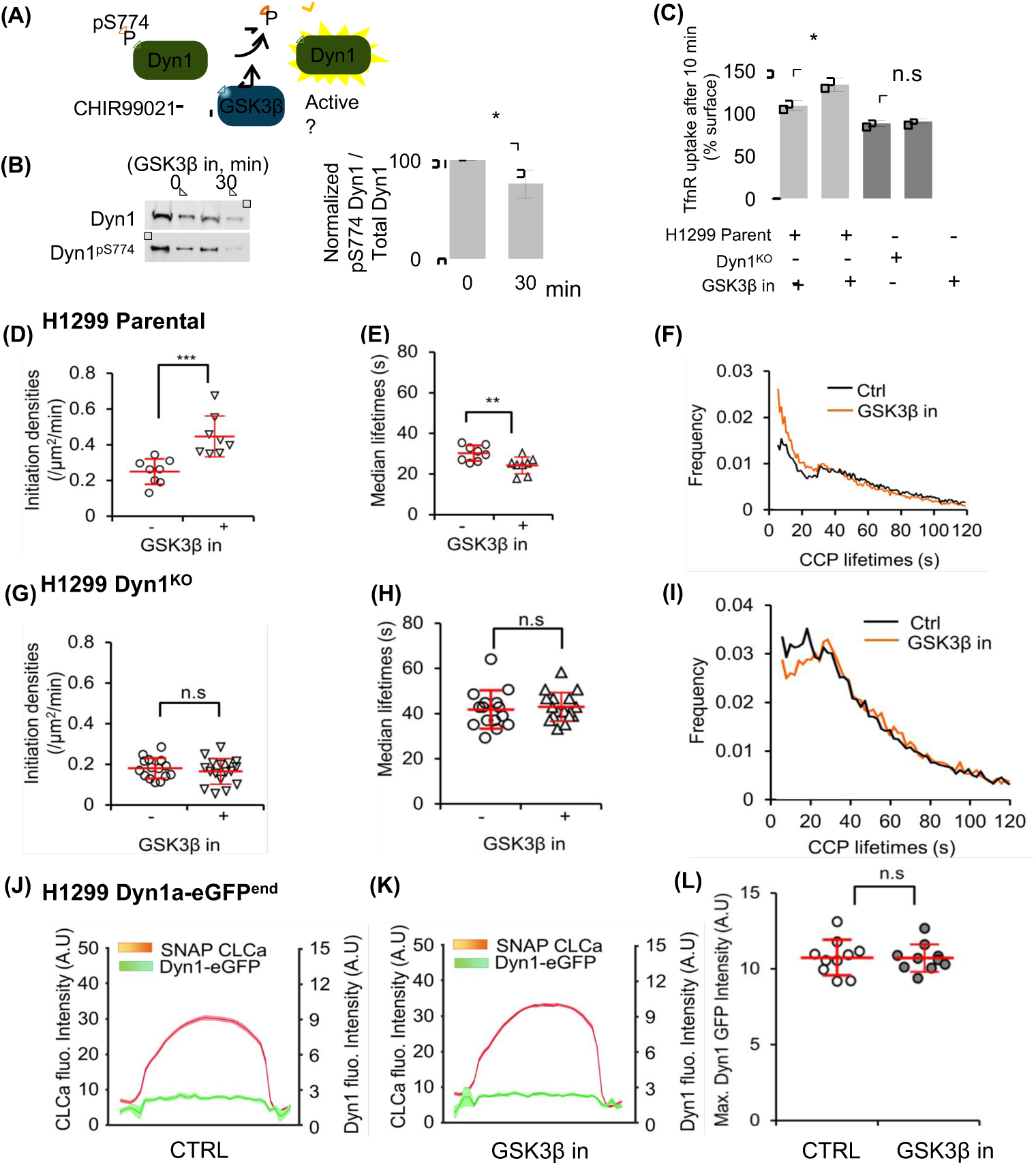
Activated Dyn1 regulates early stages of CME, even when recruited at low levels to. (A)Schematic representation of Dyn1 regulation by phosphorylation/dephosphorylation and activation upon GSK3β kinase inhibition (B) Dephosphorylation of Dyn1 S774 upon GSK3β by 20 μM CHIR99021 observed by immunoblotting using a Dyn1 phospho-specific antibody and the quantification of pDyn1/Dyn1 intensity ratios (mean ± S.D., n=3) (C) Transferrin recptor (TfnR) internalization efficiency of parental H1299 cells and Dyn1^KO^ cells and their sensitivity to GSK3β inhibition (mean ± S.D., n=3). (D) Initiation densities of bona fide CCPs and (E) their median lifetimes. Each dot represents the average value per movie, where each movie contained 1-5 cell (see Methods) (F) The distribution of CCP lifetimes measured in the absence or presence of GSK3β inhibitor. Data are derived from 10 movies each, 13346 CCPs of 40-60s lifetime analyzed from 74807 bona fide CCPs and 13494 CCPs of 40-60s lifetime were analyzed from 75426 bona fide CCPs, respectively for control and GSK3β inhibition. Similarly, the initiation densities (G), median lifetime (H) and the lifetime distribution of bona fide CCPs (I) were analyzed for H1299 Dyn1KO cells with or without GSK3β inhibition. (J,K) Average recruitment of Dyn1a eGFP^end^ to CLCa-labeled CCPs with lifetimes of 40-60s measured in the absence (K) or presence GSK3β inhibitor. (L) Maximum intensity of Dyn1a-eGFP^end^ detected at any point throughout the lifetime of an individual CCPs measured in the absence or presence of GSK3β inhibitor. (* p≤0.05, ** 01, *** p≤0.001, see Methods for description of statistical analysis used in this and other figures)

To further probe the mechanism by which activated Dyn1 accelerates CME, we introduced mRuby2 labeled CLCa into H1299 parent Dyn1^KO^ cells and measured CCP dynamics by TIRFM. Analysis of the rates of assembly and departure of CCPs revealed that GSK3β inhibition resulted in a significant increase in the rate of coated pit initiation per unit cell area (**Fig. 2D**), as well as an increase in maturation rates (i.e. decrease in lifetimes) of CCPs (**Fig. 2E**). The latter was evident in the change in lifetime distribution of all bona fide CCPs (**Fig 2F**), which displayed a more quasi-exponential profile than untreated cells, indicative of a less regulated process during early stages of CCP maturation [22]. Importantly, similar effects were observed for H1299 Dyn1a-eGFP^end^ (**Supplemental Fig S3A-C**), confirming that the C-terminally eGFP-tagged splice variant, Dyn1a, was fully functional and activated by dephosphorylation. Again, GSK3β inhibition had no effect on CCP initiation rates or lifetimes in H1299 Dyn1^KO^ cells (**Fig 2 G-I**), confirming that these changes in CCP dynamics are a result of activation of Dyn1.

We then asked whether GSK3β inhibition and activation of Dyn1 altered its recruitment to CCPs. Surprisingly, there was no significant difference in the average recruitment intensity (**Fig 2J**) of Dyn1 at CCPs. Previous studies had shown that the appearance of dynamin fluctuates at CCPs [21, 32], thus it was possible that GSK3β inhibition induces asynchronous and transient appearances of Dyn1 at CCPs that could be obscured by measuring average recruitment. Therefore, we also quantified the maximum intensity of Dyn1 recruited at any time along a CCP track. Using this orthogonal measurement, we again, saw no effect of GSK3β inhibition on Dyn1 recruitment to CCPs (**Fig 2K**). Together these data suggest that dephosphorylation and activation of Dyn1 can alter CCP dynamics and CME even when Dyn1 is present at low amounts, and that the effects of activation of Dyn1 on CCP dynamics, are not likely explained simply by its increased recruitment to CCPs.

### Sub-stoichiometric levels of Dyn1 are sufficient to stimulate CCP dynamics

It remained possible that the extremely low expression levels of Dyn1 in H1299 might limit our ability to detect GSK3β-dependent changes in its recruitment. To test this, we stably overexpressed Dyn1aWT-eGFP in H1299 Dyn1^KO^ cells at ∼20-fold levels higher than endogenous to generate Dyn1aWT-eGFP^o/x^ cells (**Fig 3A**). Importantly, overexpression of Dyn1a^WT^-eGFP, itself, did not result in any additional increase in TfnR uptake compared to the normal low endogenous levels (**Fig 3B**, see also Fig. 4G). However, as in parental and genome-edited H1299 cells, acute GSK3β inhibition in the Dyn1aWT-eGFP^O/X^ cells resulted in increased rates of TfnR uptake (**Fig 3B**) and alterations in CCP dynamics, including increased rates of CCP initiation and maturation (**Fig 3C-E**). Yet, similar to the Dyn1a-eGFP^end-^cells, GSK3β inhibition did not result in significantly enhanced recruitment of Dyn1a^WT^-eGFP to the membrane, either on average (**Fig 3F,G**) or when measured as maximum peak intensity (**Fig 3H**). Moreover, there was no evidence of a burst of Dyn1 recruitment prior to CCV formation (**Fig 3G**). Together these results suggest that the observed changes in CCP dynamics are the result of a scission-independent, early role for low levels of Dyn1 in regulating CME.

**Figure 3:**
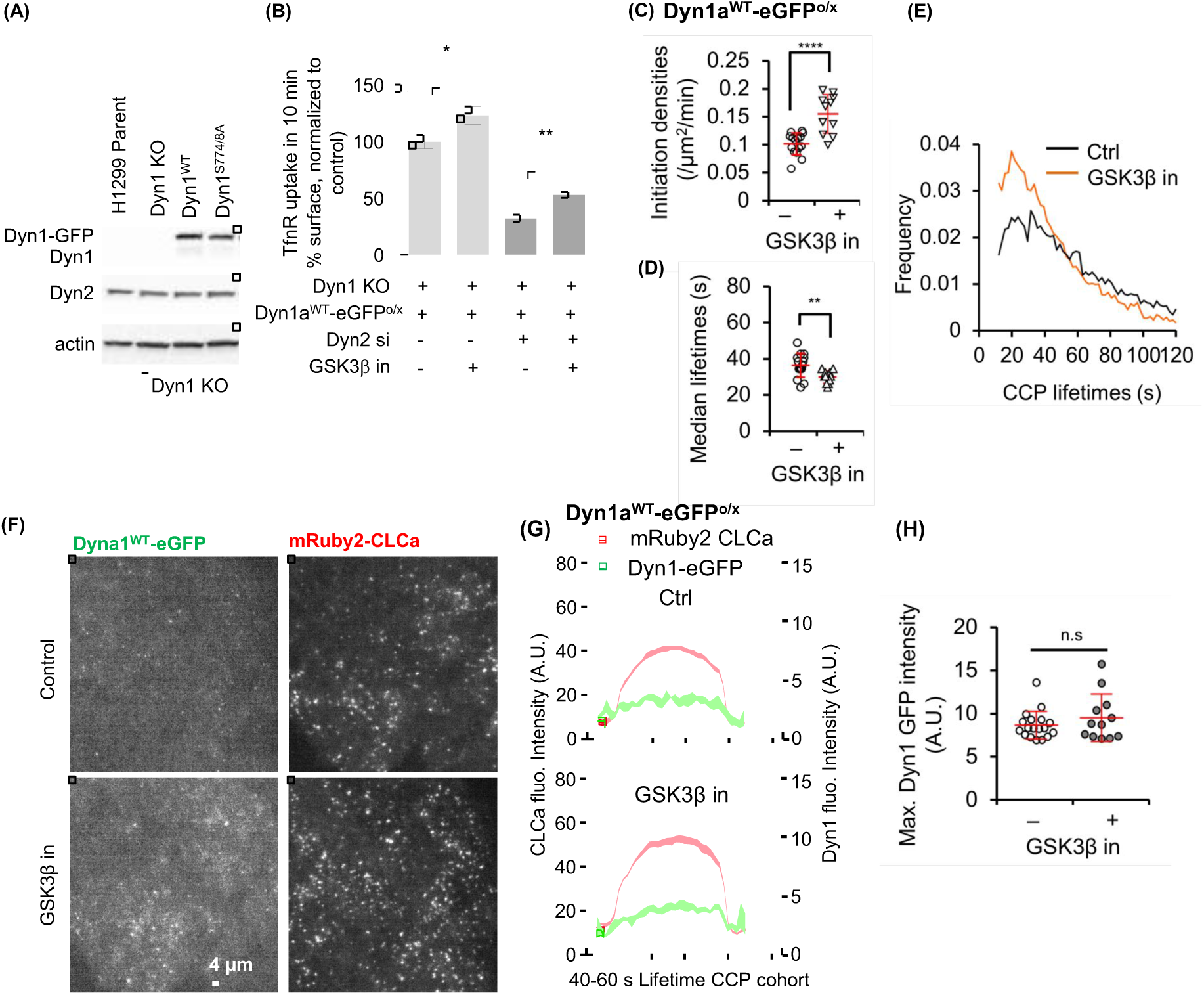
Nonphosphorylatable Dyn1 mutant mimics GSK3β β effect and can partially substitute Dyn2. Western blot showing overexpression of Dyn1^WT^-eGFP or Dyn1^S774/8A^-eGFP in Dyn1^KO^ H1299 cells.(B) Effect of siRNA knockdown of Dyn2 on TfnR internalization in Dyn1^KO^ cells reconstituted with a^WT-^eGFP and treated or not with GSK3β inhibitor. Results are normalized to rates of endocytosis in parental H1299 cells. The data represents mean ± sem of n=3 experiments containing four replicates each (*p≤0.05, ** p≤0.01). Initiation densities (C), median lifetimes (D) and the lifetime distribution (E) of bona fide CCPs analyzed in H1299 Dyn1^KO^ cells reconstituted with Dyn1^WT^-eGFP or without GSK3β inhibition, determined as in Fig. 2. (F) Representative TIRF images of overexpressed Dyn1^WT^-eGFP and mRuby2-CLCa and (G) quantification of the average recruitment of ^WT^-eGFP to CCPs, identified by mRuby2-CLCa, with lifetimes between 40-60s (14495 CCPs from a pool of 100050 bona fide Dyn1 positive CCPs from 18 movies and 9651 CCPs from a pool of 68909 bona fide CCPs from 12 movies were analyzed from control and GSK3β, respectively). (H) Maximum Dyn1a^WT^-eGFP intensity averaged among individual bona fide CCP tracks in the absence or presence of GSK3β inhibitor.

### Dephosphorylated Dynamin 1 regulates early stages of CME

Based on our finding that Dyn1 expression is required for the inhibitory effects of GSK3β on CME, we hypothesized that dephosphorylation of residues in Dyn1’s PRD should be sufficient to enhance CME efficiency. To test this, we introduced point mutations in Dyn1 at the serine residue phosphorylated by GSK3β (S774) and at the priming serine site that is responsible for recruiting GSK3β (S778). We expressed this mutant as an eGFP fusion in H1299 cells, Dyn1a^S774/8A^-eGFP, at comparable levels to Dyn1a^WT^-eGFP (**Fig 3A**). As predicted, Dyn1^S774/8A^-eGFP cells exhibited increased rates of CCP initiation (**Fig 4A**), decreased CCP lifetimes (i.e. increased rates of CCP maturation, **Fig 4B**) and changed the lifetime distribution to a quasi-exponential profile (**Fig 4C**). From these data, we conclude that dephosphorylated Dyn1 is sufficient to account for the effects of GSK3β inhibition on CCP dynamics.

**Figure 4:**
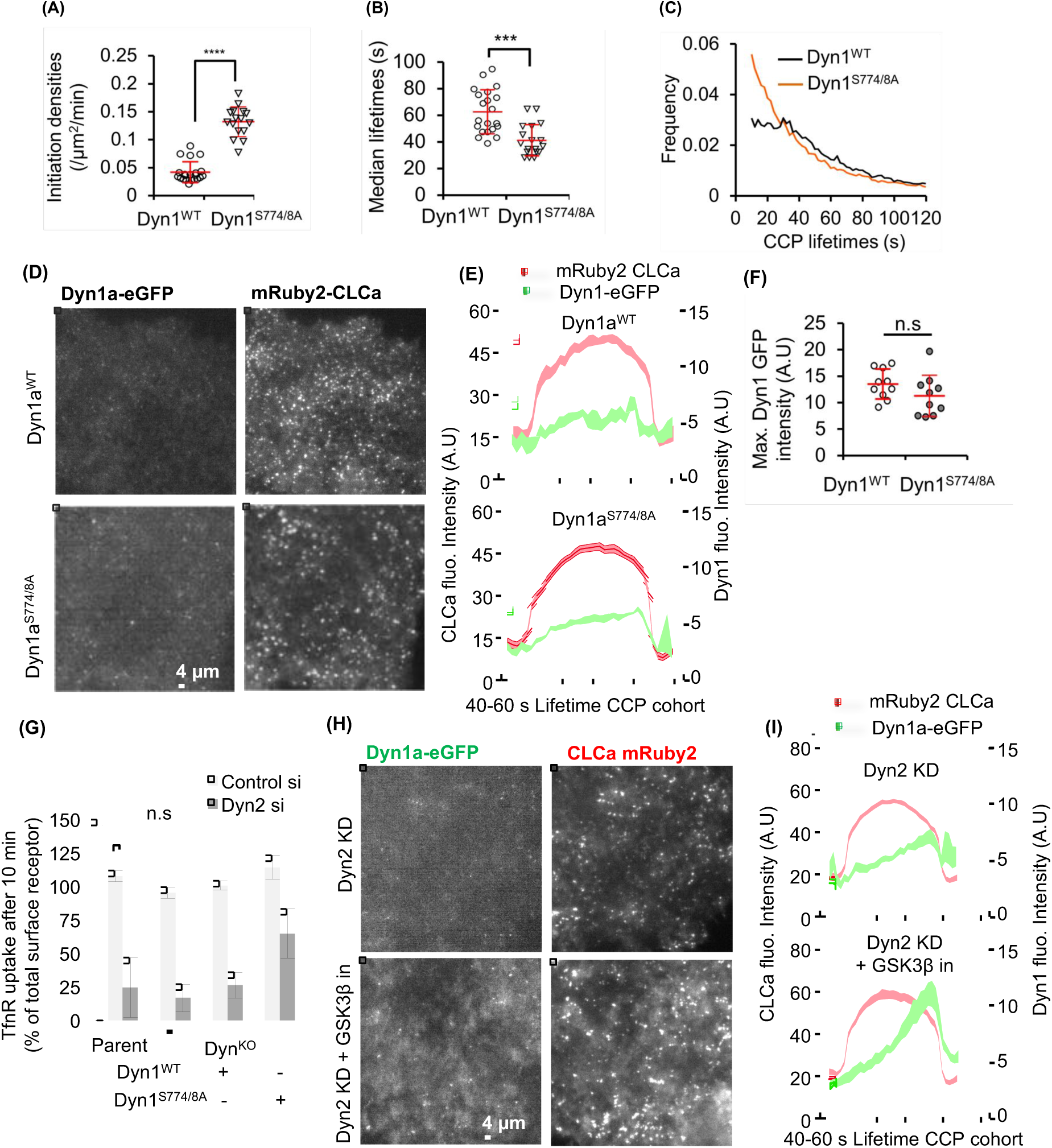
Nonphosphorylatable Dyn1 mutant mimics GSK3β effects and can partially substitute for Dyn2. Ccp initiation densities (A), median lifetimes (B) and the lifetime distribution (C) of bona fide CCPs analyzed in H1299 Dyn1^KO^ cells reconstituted with Dyn1^WT^ or Dyn1^S774/8A^-eGFP, determined as described in Fig. 2. (D) Representative TIRF images of overexpressed Dyn1^WT^-eGFP or Dyn1^S774/8A-^and mRuby2-CLCa and (E) quantification of their average recruitment to CCPs with lifetimes between 40-60s. (F) Maximum intensities of Dyn1^WT^-eGFP or Dyn1^S774/8A^-eGFP averaged among indvidual bona fide CCP tracks. (G) Effect of siRNA knockdown of Dyn2 on TfnR endocytosis in parental and Dyn1^KO^ H1299 cells and Dyn1^KO^ cells reconstituted with either Dyn1a^WT^-eGFP or Dyn1a^S774/8A-^(H) Representative TIRF images of Dyn2 siRNA-treated Dyn1^KO^ cells overexpressing Dyn1a^WT-^and mRuby2-CLCa treated or not with GSK3β inhibitor and (I) quantification of the average recruitment of Dyn1^WT^-eGFP to CCPs with lifetimes between 40-60s in Dyn2 knockdown cells treated or not GSK3β inhibitor.

Surprisingly, even the nonphosphorylatable Dyn1a^S774/8A^-eGFP mutant was not efficiently recruited to CCPs and failed to display a pronounced late burst of recruitment accompanying membrane scission (**Fig 4D-F**). Interestingly, the changes in CCP dynamics in Dyn1a^S774/8A-^eGFP expressing cells were not reflected in significantly increased rates of TfnR uptake, presumably due to compensatory changes that occur upon prolonged expression of activated Dyn1 vs. acute activation (**Fig. 4G)**. However, unlike parental H1299 cells or Dyn1a^WT^-eGFP cells, Dyn1^KO^ cells reconstituted with Dyn1a^S774/8A^-eGFP exhibited significant residual levels of TfnR uptake upon siRNA knockdown of Dyn2 (**Fig 4G**), consistent with functional activation of Dyn1. Moreover, upon siRNA knockdown of Dyn2, even Dyn1a^WT^-eGFP tended to exhibit a burst of recruitment prior to CCV formation (**Fig 4H,I)**, suggesting its activation as part of a compensatory mechanism to restore CME [23]. Under these conditions, GSK3β inhibition appears to enhance the burst of Dyn1a^WT^-eGFP recruitment prior to scission (**Fig 4H,I**).

From these data, we conclude that Dyn1 is negatively regulated in non-neuronal cells through GSK3β-dependent phosphorylation of S774, and that dephosphorylated, active Dyn1 regulates early stages of CME even when present at low (nearly undetectable, in the case of parental H1299 cells) levels on CCPs. Importantly, overexpressed Dyn1, even when activated by mutation or GSK3β inhibition (**Fig. 3B**) does not fully compensate for loss of Dyn2 function in CME, hence both isoforms have partially divergent functions.

### A549 cells express high levels of Dyn1 that can partially substitute for Dyn2

We previously reported that several lung cancer cell lines express high levels of Dyn1 [35, 39]. For example, A549 non-small cell lung cancer cells express∼5-fold higher levels of Dyn1 than Dyn2 [39], corresponding to ∼20-fold higher levels of Dyn1 than in H1299 cells (**Supplemental Fig S4A**). Reflective of these high levels of Dyn1 expression, siRNA knockdown of both Dyn1 and Dyn2 is necessary for potent inhibition of TfnR uptake in A549 cells (**Supplemental Fig S4B**). Therefore, we reasoned that it might be possible to individually knockout Dyn1 and Dyn2 in A549 cell lines for reconstitution experiments. Thus, we used CRISPR/Cas9n to generate a complete knockout of Dyn1 (Dyn1^KO^) or Dyn2 (Dyn2^KO^) in A549 cells (**Fig. 5A, Supplemental Fig S4C**) and then introduced mRuby-CLCa to track CCP dynamics. Acute inhibition of GSK3β had no effect on the rates of CCP initiation or maturation in Dyn1^KO^ A549 cells, but significantly stimulated the rate of CCP initiation and decreased the lifetimes of CCPs in Dyn2^KO^ A549 cells (**Fig 5B,C**). These data show that the two isoforms differentially regulate early stages of CME and confirm that the effects of GSK3β inhibition on CME depend on Dyn1, but not Dyn2.

**Figure 5:**
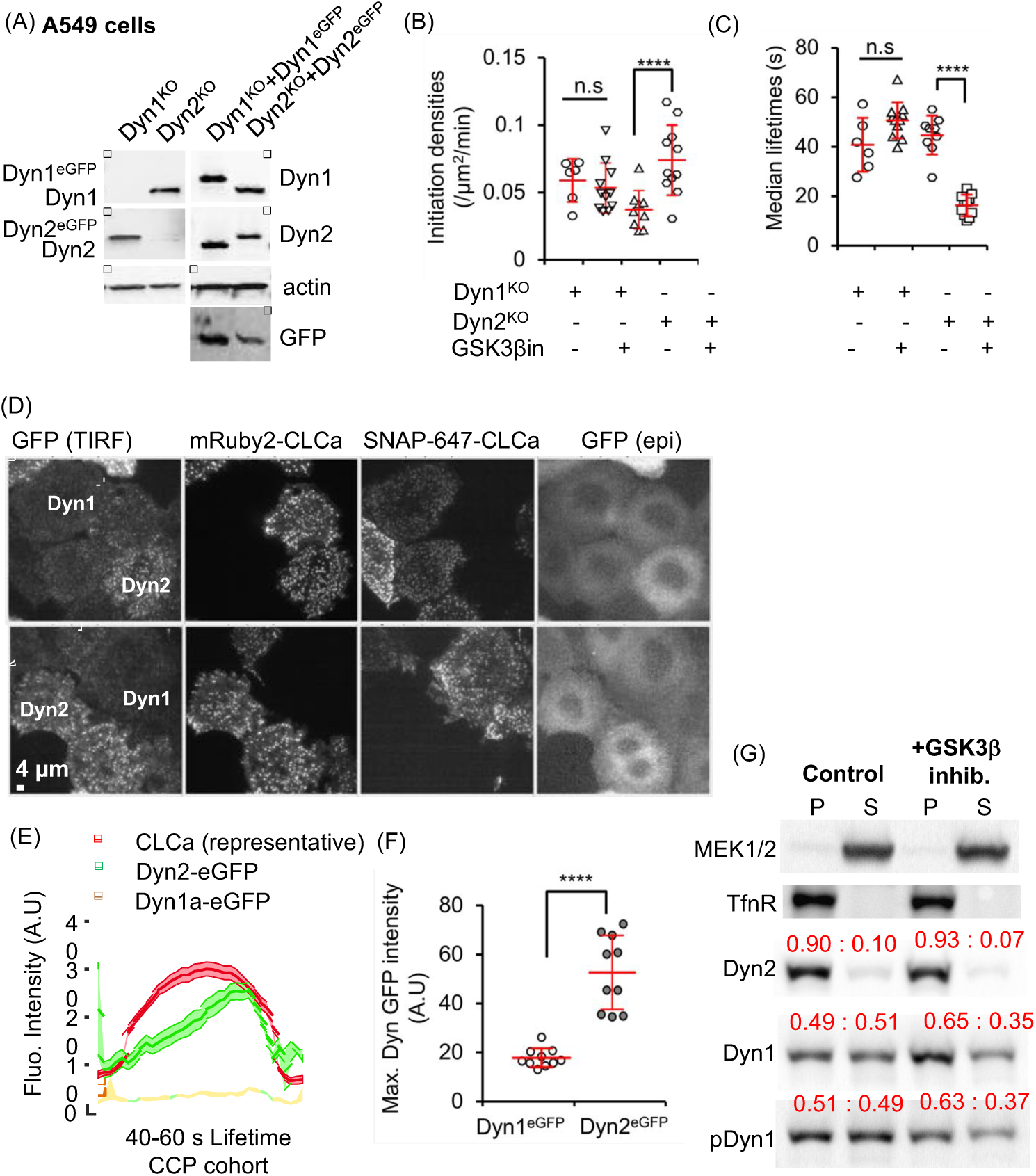
Dyn1 and Dyn2 are differentially recruited to CCPs and differentially required for GSK3β β-regulated CME. Immunoblot validation of Dyn1 and Dyn2 KO A549 cells and their corresponding reconstitution at near endogenous levels with eGFP-labelled Dyn1 or Dyn2. GFP blot shows that in A549 cells Dyn1 is expressed at ∼5-fold higher levels than Dyn2. CCP initiation densities (B), and median lifetimes (C) in Dyn1 or Dyn2 knockout cells with or without GSK3β inhibition, determined as described in Fig. 1. (D) Representative TIRF and epifluorescent (epi) images of co-cultured and Dyn1^KO^(Dyn1a-eGFP:SNAP-CLCa) Dyn2^KO^(Dyn2-eGFP:mRuby2-CLCa) cells allowing direct comparison of Dyn1a-eGFP vs Dyn2-^eGFP^ recruitment to CCPs in A549 cells. (E) Quantification of the average recruitment of Dyn1a-eGFP 2-eGFP to CCPs with lifetimes between 40-60s (4420 CCPs from a pool of 12555 Dyn1a-eGFP positive CCPs and 3961 CCPs from a pool of 12766 Dyn2-eGFP positive CCPs from a total of 11 movies were identified to have a lifetime between 40-60s). Data are obtained from cells co-imaged either for SNAP(647)-CLCa (and Dyn1a-eGFP) or mRuby-CLCa (and Dyn2-eGFP). (F) Maximum intensities of Dyn1a-eGFP or Dyn2-eGFP averaged among individual bona fide CCP tracks. (G) subcellular fractionation of parental A549 cells into membrane (P) vs cytosolic (S) fractions and western blotted for the indicated proteins. Cytosolic MEK1/2 and membrane-associated TfnR serve as controls for fractionation. Quantification is shown in red above each band as the fraction of total protein in the P vs S fraction. Results are representative of 3 independent experiments.

To directly and quantitatively compare the relative recruitment efficiencies of the two isoforms to CCPs, we reconstituted these knockout cells with their respective eGFP-tagged isoforms and sorted for expression comparable to their endogenous levels (i.e. in these A549 cells we chose cells in which Dyn1a-eGFP levels were ∼5-fold higher than Dyn2-eGFP) (**Fig 5A**). Additionally, we introduced SNAP-CLCa and mRuby2-CLCa in Dyn1a-eGFP and Dyn2-eGFP cells, respectively so that we could distinguish the two A549 cell lines (i.e. Dyn1^KO^:Dyn1aeGFP:SNAP-CLCa from Dyn2^KO^:Dyn2-eGFP:mRuby2-CLCa) while imaging them in the same TIRF field of view, under the same conditions (**Fig 5D**). These data directly show the differential recruitment efficiencies of Dyn1 and Dyn2 to CCPs. Live cell imaging revealed the typical gradual accumulation and burst of Dyn2-eGFP recruitment to CCPs when averaged over the cohort of 40-60s lifetime CCPs (**Fig 5E**). However, under identical imaging conditions of the same fluorophore, Dyn1a-eGFP was recruited, on average at least 10-fold less efficiently, even though it is expressed at higher abundance. The maximum intensity of tagged Dyn2 vs Dyn1 recruitment was also higher, albeit showing only an ∼3-fold differential (**Fig 5F**). A likely explanation for the differences in average and peak measurements is that in A459 cells, Dyn1aeGFP does display a slight burst of recruitment at late stages of CCV formation that is visible when the Dyn1 signal is rescaled (**Supplemental Fig S4D**).

Finally, to verify our results using an independent method, we performed Western blotting after subcellular fractionation and isolation of membrane vs. cytosolic fractions, as verified using membrane-associated TfnR and cytosolic MEK1/2 as markers (**Fig 5G**). Under these fractionation conditions, ∼90% of Dyn2 is membrane associated, whereas only 50% of Dyn1 sediments with the membrane fraction (**Fig 5G**). We observed a consistent, ∼20% increase of membrane-associated Dyn1 upon GSK3β inhibition that was not detected by TIRFM. These biochemical data indicate a greater extent of membrane association of both active and inactive Dyn1 than detected at CCPs by TIRFM. The differences could reflect recruitment of Dyn1 to sites on the plasma membrane other than CCPs, as has been previously reported [40]. The ∼20% increase in recruitment of activated Dyn1 likely reflects the increase in number of CCPs that occurs upon GSK3β inhibition, rather than an increase in Dyn1 per CCP. Consistent with TIRFM data, the distribution of phosphorylated Dyn1 (detected with an S774 phospho-specific antibody) was indistinguishable from total Dyn1 (i.e. there was no de-enrichment of phosphorylated Dyn1 in the membrane-bound fractions). These data confirmed that dephosphorylation of Dyn1 on S774 by GSK3β inhibition does not enhance its recruitment to CCPs. Thus, the effects of activated Dyn1 on CCP initiation and maturation occur either independently of its direct association with CCPs, or, more likely, are manifested by very low levels of CCP-associated dephosphorylated Dyn1.

### Dyn1 and Dyn2 do not efficiently co-assemble

Dynamin exists as a tetramer in solution [41, 42] and assembles into higher order helical oligomers on the membrane. Exploiting Dyn1^KO^ and Dyn2^KO^ A549 cells reconstituted with Dyn1a‐ or Dyn2-eGFP, respectively, we next assessed the degree to which Dyn1 and Dyn2 form hetero-tetramers in solution. Dyn1‐ or Dyn2-eGFP were efficiently immunoprecipitated with anti-eGFP nanobodies and the immunobeads were washed with 300 mM salt to disrupt any potential higher order dynamin assemblies before measuring the fraction of Dyn2 or Dyn1 that co-precipitated. Under these conditions we pulled down nearly 100% of the eGFP-tagged dynamins, but only ∼30% of Dyn2 with Dyn1-eGFP and <5% of Dyn1 with Dyn2-eGFP (**Supplemental Fig S5A).** The difference in the extent of hetero-tetramerization is consistent with the ∼5-fold higher levels of expression of Dyn1 vs Dyn2 in these cells. Thus, the two isoforms predominantly exist as homo-tetramers in solution.

We also examined the relative abilities of Dyn1 and Dyn2 to co-assemble into higher order structures *in vitro.* For this we used a dominant-negative Dyn1 mutant (Dyn1^S45N^) defective in GTPase activity, which when co-assembled with wild-type dynamin into higher order oligomers ‐on lipid nanotubes will inhibit total assembly-stimulated GTPase activity through the intercalation of GTPase-defective subunits adjacent to wild-type subunits [43, 44]. As expected, Dyn1^S45N^ efficiently co-assembles with Dyn1^WT^ such that when present at equimolar levels the total assembly-stimulated GTPase activity is inhibited by 50%. In contrast, at the same concentrations of Dyn1^S45N^, Dyn2 GTPase activity was significantly less affected (**Supplemental Fig S5B)**, indicating that Dyn2 less efficiently co-assembles into higher order oligomers with the mutant Dyn1. Thus, consistent with their differential recruitment to CCPs, even when present at comparable levels of expression in the same cell type, the two isoforms only weakly interact.

### Genome edited cells reveal that Dyn1 and Dyn2 are recruited to most CCPs in A549 cells

Our results establish that Dyn1 and Dyn2 are differentially recruited to CCPs in non-neuronal cells, and that, on average, Dyn1 is recruited at much lower levels than Dyn2. Despite this, acute activation of Dyn1 globally alters CCP dynamics. Thus, we next directly compared the recruitment of Dyn1 and Dyn2 to CCPs to determine whether Dyn1 is recruited at low levels to all CCPs or instead might be recruited at higher levels to a subpopulation of CCPs. Such heterogeneity would be lost by averaging. For this, we took advantage of the higher levels of Dyn1 expression in A549 cells and generated double genome-edited cells expressing Dyn1aeGFP and Dyn2-mRuby. We first used ZFNs to generate Dyn2 mRuby2-edited A549 cells, and subsequently introducing a C-terminal eGFP to the Dyn1a splice variant using CRISPR/Cas9, as described earlier (**Fig. 1A**, see Methods). This yielded an A549 cell line homozygous for endogenously-tagged Dyn2-mRuby2 and heterozygous for endogenously-tagged Dyn1a-eGFP (2 of 3 Dyn1 alleles tagged in these triploid A549 cells) (**Fig 6A**). We confirmed that the double genome-edited cells exhibited comparable rates of TfnR uptake, as well as the degree of dependence on Dyn2 for CME, relative to the parent cells (**Fig 6B**). SNAP-CLCa was introduced into these cells by lentiviral transfection (**Fig. 6C)** and we confirmed that GSK3β inhibition resulted in increased rates of CCP initiation, reduced CCP lifetimes and altered the lifetime distributions of CCPs (**Fig 6D-F)**, as in the parental cells. Thus, the genome-edited Dyn isoforms were functionally active.

**Figure 6:**
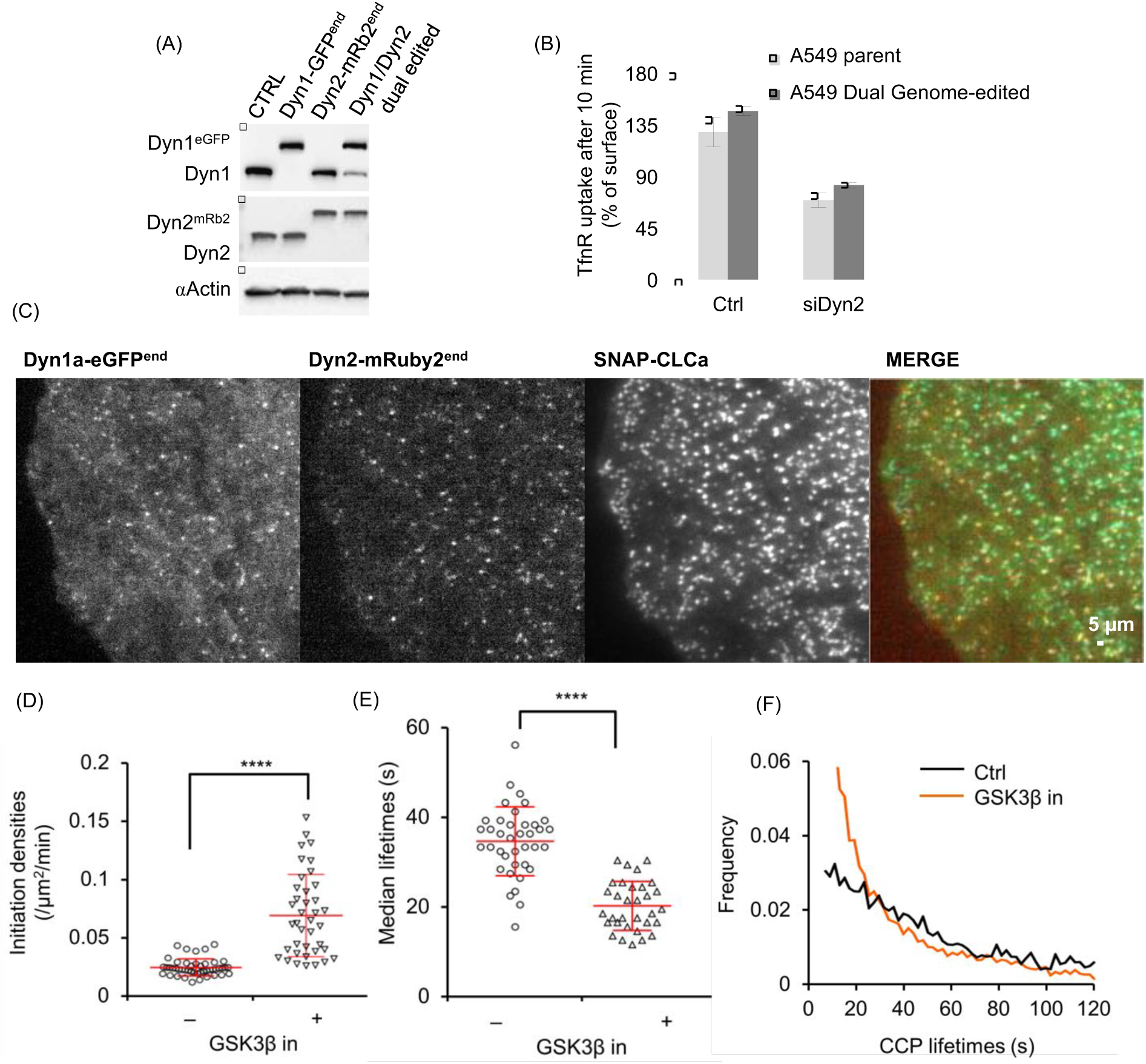
Generation and characterization of dual genome-edited Dyn1a-eGFP and Dyn2-mRuby A549 cells. (A)Immunoblot validation of Dyn1a-eGFP and Dyn2-mRuby2 single‐ and dual-genome edited A549. (B) TfnR endocytosis in dual genome edited A549 cells compared to parental A549 cells and their sensitivity to siRNA-mediated Dyn2 knockdown. (C) Representative TIRF images of Dyn1 and Dyn2 distribution relative to CLCa in dual genome-edited A549 cells. CCP initiation densities (D), median lifetimes (E) and the lifetime distribution (F) of bona fide CCPs in dual genome-edited A549 cells with or without GSK3β inhibition, determined as in Fig. 2.

We next assessed the interplay between Dyn1a-eGFP and Dyn2-mRuby using three-color live-cell TIRFM imaging at 0.5 Hz (2s/frame) (**Fig 7A, Supplemental Movie S3**). As reported previously, we detected fluctuations of both Dyn1 and Dyn2 at CCPs over their lifetimes (examples shown in **Fig 7B**), and frequently detected a burst of Dyn2 just prior to CCV formation. In many cases we also detected a burst of Dyn1 recruitment, albeit to a lesser degree. For more quantitative analysis of these data, we applied the 3-channel functionality of our cmeAnalysis package to perform three-color master/slave analyses [22]. Using clathrin as the ‘master’ channel and Dyn1 and 2 as ‘slave’ channels, we determined whether the clathrin tracks contained either Dyn1, Dyn2, both or neither. Individual CCP tracks were considered positive for Dyn1 and/or Dyn2 if the intensities of Dyn1/2 signals detected at the position of the clathrin tag were significantly higher than the local Dyn1/2 background signal around the clathrin tag position for a period of time exceeding random associations, as previously described [22]. This analysis revealed that in double genome-edited Dyn1a-eGFP^end^/Dyn2-mRuby^end^ A549 cells both Dyn2 and Dyn1 could be robustly detected in ∼75% of all bona fide CCPs (**Fig 8A**). Moreover, in this population of CCPs a clear burst of recruitment of both Dyn1a-eGFP and Dyn2-mRuby could be detected prior to CCV formation. Importantly, the apparently higher levels of recruitment of Dyn1-eGFP vs Dyn2-mRuby in these genome-edited cells is not a reflection of protein levels, but rather of imaging conditions and brightness for two different fluorophors (compare with **Fig 5E**). The remaining CCPs were roughly equally distributed as Dyn1 only, Dyn2 only and both Dyn1 and Dyn2 negative subpopulations (**Fig. 8A**). Note that the Dyn2 levels in the “Dyn1 only” CCPs were still on average higher than background (Dyn1/Dyn2 negative), reflecting the stringency of our master/slave detection, and suggesting that Dyn2 is recruited to >90% of all CCPs, albeit to variable extents.

**Figure 7:**
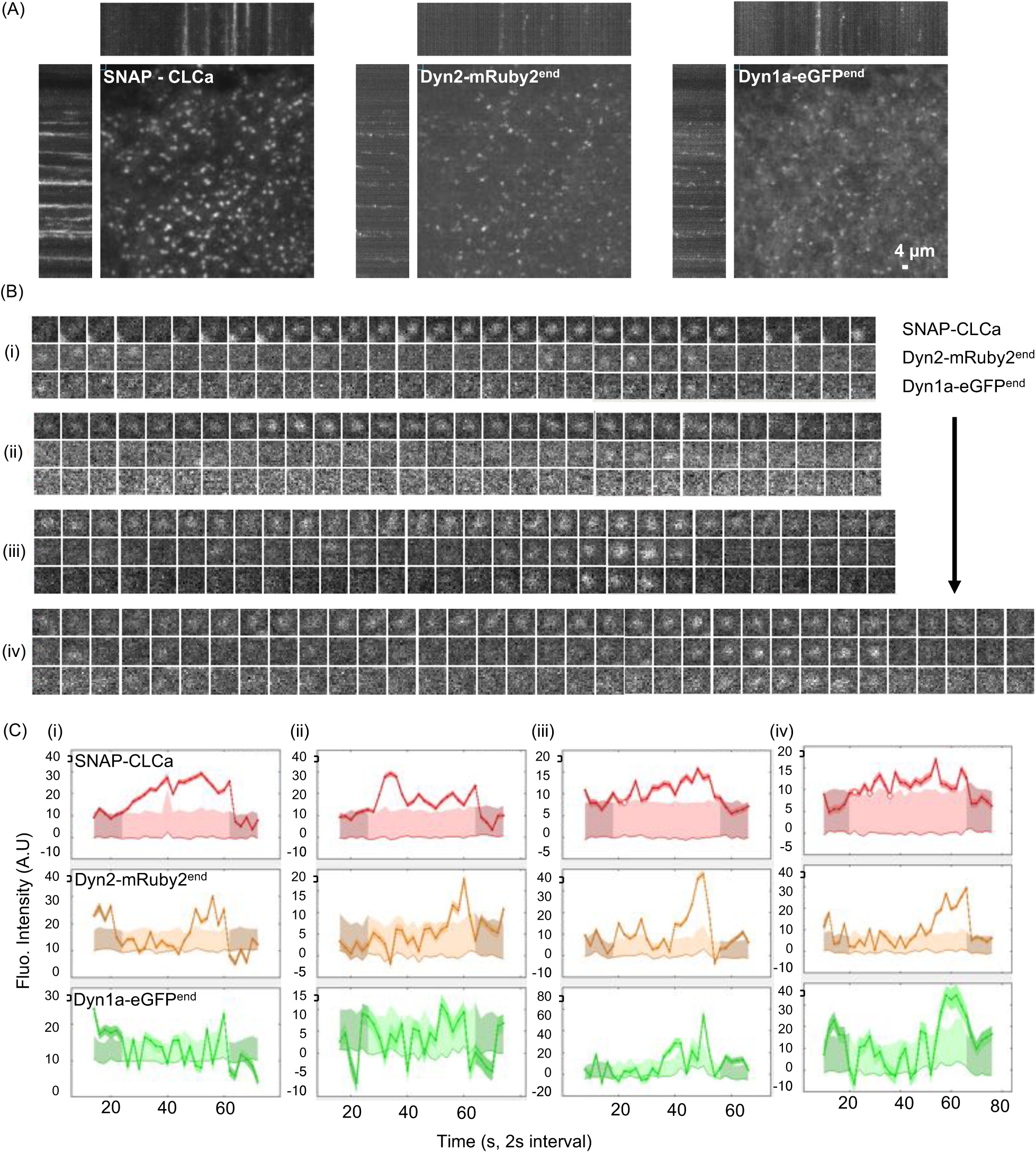
Tracking clathrin and dynamins in dual genome-edited Dyn1a-eGFP and Dyn2-mRubyA549 cells. (A)Representative TIRF images and corresponding kymographs of dynamic behavior of overexpressed SNAP-CLCa, Dyn2-mRuby^end^, and Dyn1a-eGFP^end^ in dual genome-edited A549 cells. See Supplemental Movie S3. (B) Examples of Dyn1 and Dyn2 dynamics at individual CCPs (i-iv) and (C) their corresponding quantitative traces.

**Figure 8:**
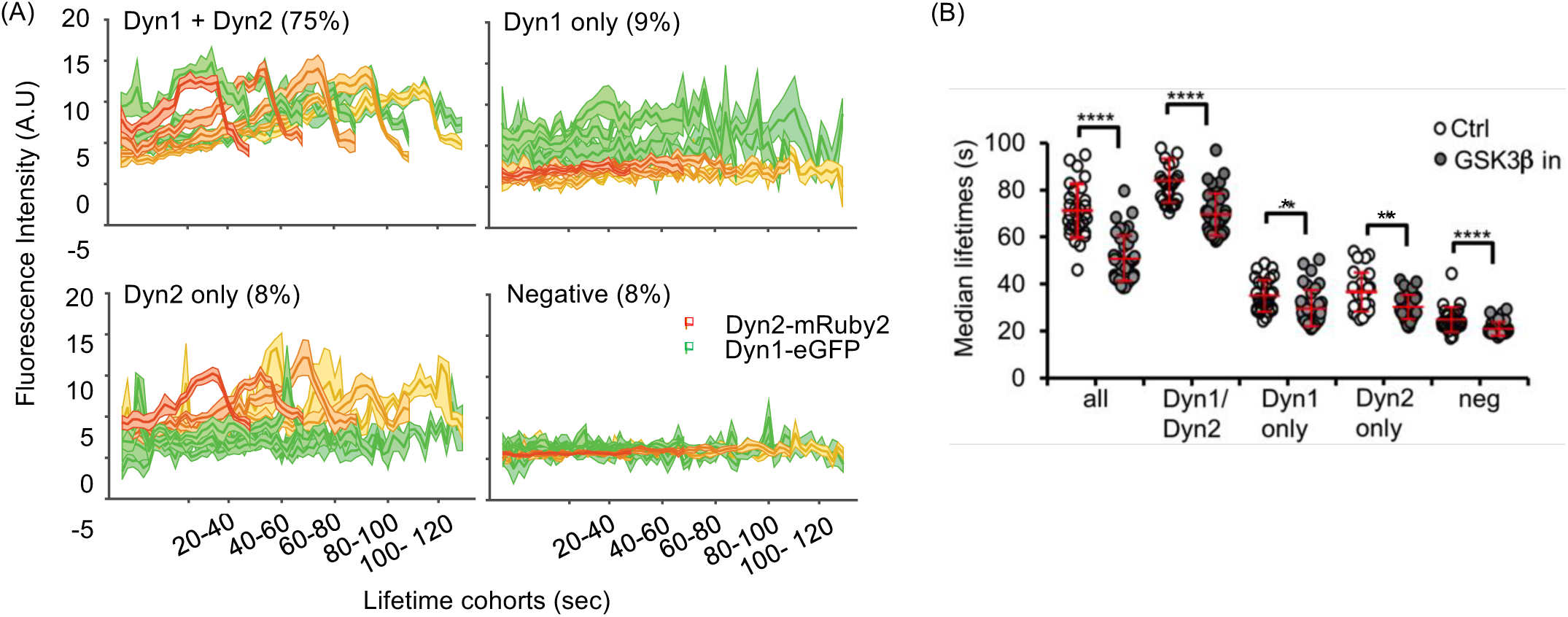
Dyn1 and Dyn2 are recruited to same CCPs and Dyn1 activation alters the dynamics of all CCP subpopulations. (A) Triple color, master/slave analyses of average dynamics of recruitment of Dyn2-mRuby^endo^ and/or Dyn1a-eGFP^endo^ to lifetime cohorts of SNAP-CLCa labeled CCPs identifies Dyn1/Dyn2 positive, Dyn1 only, Dyn2 only and Dyn1/2 negative subpopulations of CCPs. The percentage of detected CCPs in each class indicated. (B) Effect of GSK3β inhibition on the median lifetimes of compositionally-distinct CCP subpopulations.

We next compared per-cell median lifetimes of CCPs relative to their dynamin isoform composition and found that CCPs bearing higher levels of Dyn2 and Dyn1 exhibited longer lifetimes (median ∼80s) than single-positive CCPs (median ∼38s) (**Fig. 8B**). CCPs that failed to detectably recruit either isoform were the shortest lived (median ∼20s). These findings are consistent with previous data suggesting that a threshold level of Dyn2 recruitment is required for efficient CCP maturation [22, 34]. All of these CCP subpopulations, with the exception of the Dyn2 only CCPs, which nonetheless trended downwards, showed a significant decrease in CCP lifetimes upon inhibition of GSK3, consistent with other data that only low levels of Dyn1 are required to alter CCP maturation.

### SNX9 is a required for activated Dyn1-dependent effects on CCP maturation

Our findings thus far point to isoform-specific functions of Dyn1 and Dyn2 and hence suggest the existence of isoform-specific binding partners. Dyn1 and Dyn2 are >80% identical except for their C-terminal PRDs, which are only 50% identical and likely determine isoform-specific interactions with SH3 domain-containing proteins. The Dyn1^KO^ and Dyn2^KO^ A549 cells provide an opportunity to measure Dyn2 and Dyn1-dependent CME, respectively, without the possibility of compensation. Thus, we measured, by TfnR uptake, the effects of siRNA knockdown of several known SH3 domain containing binding partners on Dyn2-dependent CME in the Dyn1^KO^ cells, and on Dyn1-dependent CME in the Dyn2^KO^ cells. Knockdown of these dynamin partners has only mild effects on TfnR uptake in parental A549 cells and in Dyn1^KO^ cells, whose endocytosis is exclusively Dyn2-dependent (**Fig. 9A**). Whether these mild effects reflect partial redundancy with other dynamin partners, activation of compensatory mechanisms [23], or that these factors, which were identified primarily as dynamin partners in brain lysates, play only minor roles in TfnR uptake in non-neuronal cells, cannot be discerned from these studies. Interestingly, siRNA knockdown of Grb2 appeared to inhibit TfnR uptake in Dyn1^KO^ cells by ∼20%, while not affecting TfnR uptake in either parental or Dyn2^KO^ cells. This suggests that Grb2 might preferentially function together with Dyn2 in CME, and that its depletion in parental cells can be compensated for by Grb2-independent Dyn1 activity. In contrast, siRNA knockdown of SNX9 only mildly inhibited Dyn2-dependent TfnR uptake in parental A549 and had no significant effect on TfnR uptake in Dyn1^KO^ cells, but decreased TfnR uptake in Dyn2^KO^ cells by >50% (**Fig 9A**). Thus Dyn1-dependent endocytosis appears to be particularly sensitive to SNX9 knockdown.

**Figure 9:**
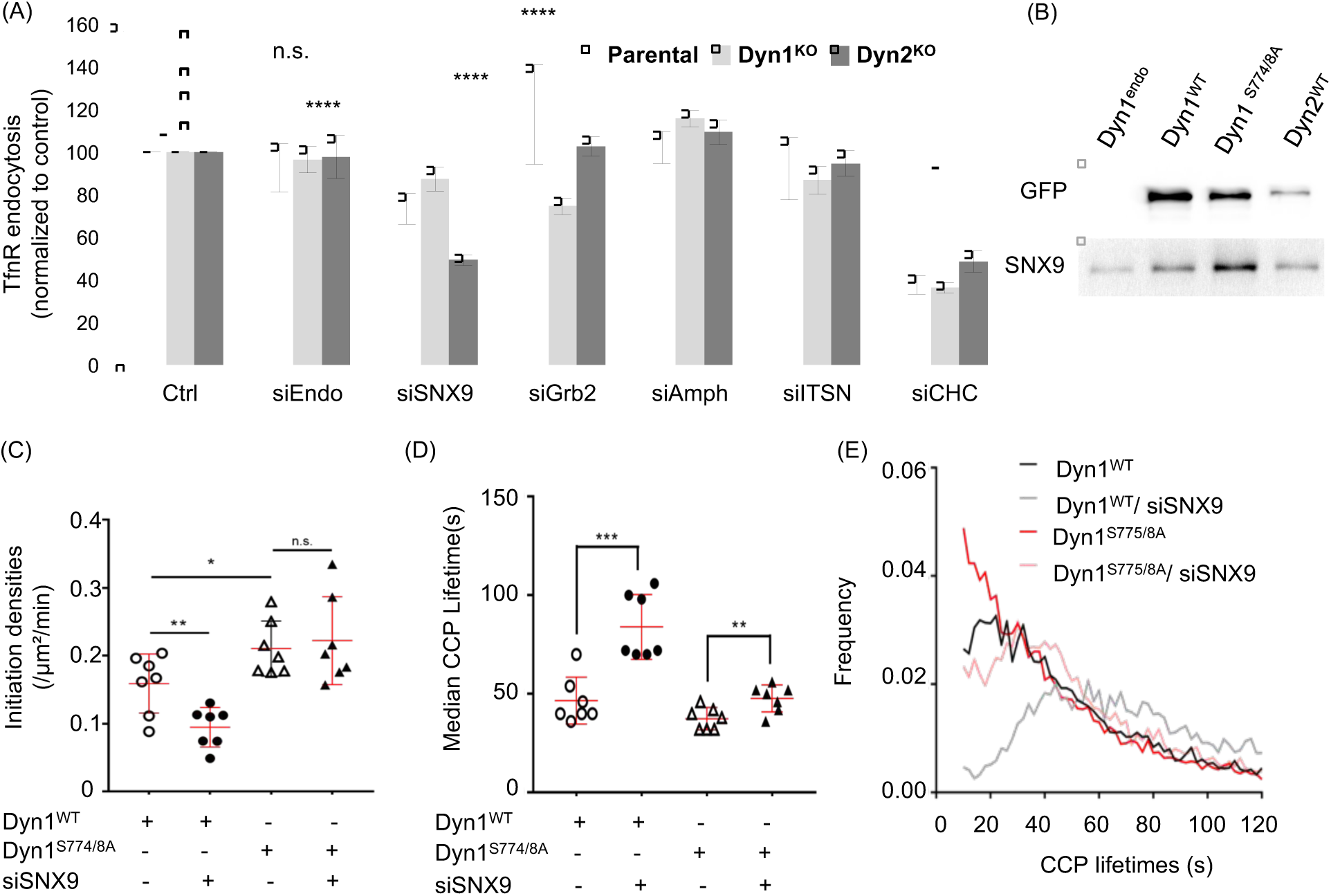
SNX9 preferentially binds activated Dyn1 and is required for Dyn1-dependent changes lifetime distribution of CCPs. Effect of siRNA knockdown of the indicated dynamin SH3 domain-containing endocytic accessory proteins on TfnR endocytosis in parental, Dyn1^KO^ and Dyn2^KO^ A549 cells. siEndo refers to siRNA knockdown of endophilin A1,2 and 3, siITSN refers to siRNA knockdown of intersectins 1 and 2, all others were single siRNAs. Knockdown efficiencies were determined to be >85% by western blotting. Data are normalized to the extent of TfnR uptake in control siRNA-treated parental, Dyn1^KO^ and Dyn2^KO^ which is set to 100, to allow direct comparison of the relative effects of siRNA knockdowns. (B) pulldown of Dyn1a^WT^-eGFP, Dyn1a^S774/8A^-eGFP or Dyn2^WT^-eGFP expressed in Dyn1^KO^ or ^KO^ A549 cells, respectively using anti-eGFP nAbs. Parental cells that do not express an eGFP-tagged protein are used as control. The pulldown fractions were analyzed by immunoblot. Effect of siRNA-mediated knockdown on (C) CCP initiation densities, (D) median lifetimes and (E) lifetime istribution of bona fide CCPs in Dyn1^KO^ H1299 cells overexpressing either Dyn1a^WT^-eGFP or ^S774/8A^-eGFP (data are derived from 7 movies for each condition with each movie consisting of 1-3 cells. Each data point is the average value from a single movie. (* p≤ 0.05, ** p≤ 0.01, *** p≤ 0.001)

We next tested whether SNX9 preferentially interacts with Dyn1 vs Dyn2 by GFP pulldown assays using Dyn1^KO^ and Dyn2^KO^ A549 cells reconstituted with either Dyn1a^WT^- or Dyn1^S774/8A^-eGFP or with Dyn2-eGFP, respectively. Consistent with previous results [45, 46], we confirmed that SNX9 binds both Dyn1 and Dyn2 (**Fig 9B**). However, the ratio of SNX9 binding to Dyn1 vs Dyn2 was 1.7±0.6 (mean ± sem, n=3), indicative of a slight preference for Dyn1. Importantly, SNX9 showed a marked preference for binding to the nonphosphorylated and active Dyn1^S774/8A^-eGFP. The ratio of SNX9 binding to Dyn1^S774/8A^ vs Dyn1^WT^ was 3.6±0.9 (mean ± sem, n=3). These data suggested that SNX9 might be a preferential functional partner of activated Dyn1.

We next asked whether SNX9-Dyn1 interactions were required for the effects of activated Dyn1 on CCP initiation rates, CCP maturation or both. For this we returned to the Dyn1^KO^ H1299 cells reconstituted with Dyn1^WT^ vs Dyn1^S774/8A^ and tested whether the selective effects of Dyn1^S774/8A^ on CCP dynamics (**Fig. 4A-C**) were dependent of SNX9. Knockdown of SNX9 decreased the rate of CCP initiation in Dyn1^WT^, but was not required for the enhanced rate of CCP initiation triggered by Dyn1^S774/8A^ expression (**Fig. 9C**). Thus, other, yet unidentified binding partners are responsible for the Dyn1-dependent effect on CCP initiation. SNX9 knockdown also led to an increase in the median CCP lifetimes in both Dyn1^WT^ and Dyn1^S774/8A^ expressing cells (**Fig. 9D**). These data suggest that SNX9 functions in both Dyn1-dependent and independent stages of CCP maturation. Consistent with this, SNX9 knockdown also abrogated the effects of Dyn1^S774/8A^ expression on the lifetime distribution of bona fide CCPs (**Fig. 9E**), reverting the quasi-exponential distribution seen in Dyn1^S774/8A^ to a distribution nearer to control. The strong effect of SNX9 knockdown is also seen in the rightward shift of the lifetime distribution of Dyn1^WT^ cells treated with SNX9 siRNA. Together these data suggest multiple roles of SNX9 at multiple stages of CME, including the support of Dyn1’s early functions in accelerating CCP maturation.

### Dyn1 is activated downstream of the EGFR

We have shown that strong pharmacological inhibition of GSK3β activates Dyn1 in non-neuronal cells and results in increased rates of CCP initiation and maturation, leading to increased rates of TfnR uptake via CME. However, it is not clear whether this regulatory effect on Dyn1 function modulates CME under more physiologically-relevant conditions. To test this, we treated serum-starved A549 cells with EGF, which is known to activate Akt and in turn to phosphorylate and inactivate GSK3β [47]. We confirmed that GSK3β is phosphorylated in EGF-treated cells and that this resulted in reduced levels of phosphorylation of Dyn1 at S774 (**Fig 10A**, quantified in **Fig 10B,C**). As predicted by the results of inhibitor experiments, EGF treatment of serum-starved cells also increased the rate of CCP initiation (**Fig 10D**), decreased CCP lifetimes (**Fig 10E**) and, compared to control cells, resulted in a shift of the lifetime distributions of bona fide CCPs to a more quasi-exponential distribution (**Fig 10F**). Importantly, the effects of EGF treatment on CCP initiation rate and lifetimes were not seen in A549 Dyn1^KO^ cells (**Fig 10G,H**). These data suggest that Dyn1 can be activated to alter CCP dynamics under physiological conditions through signaling downstream of EGFR.

**Figure 10:**
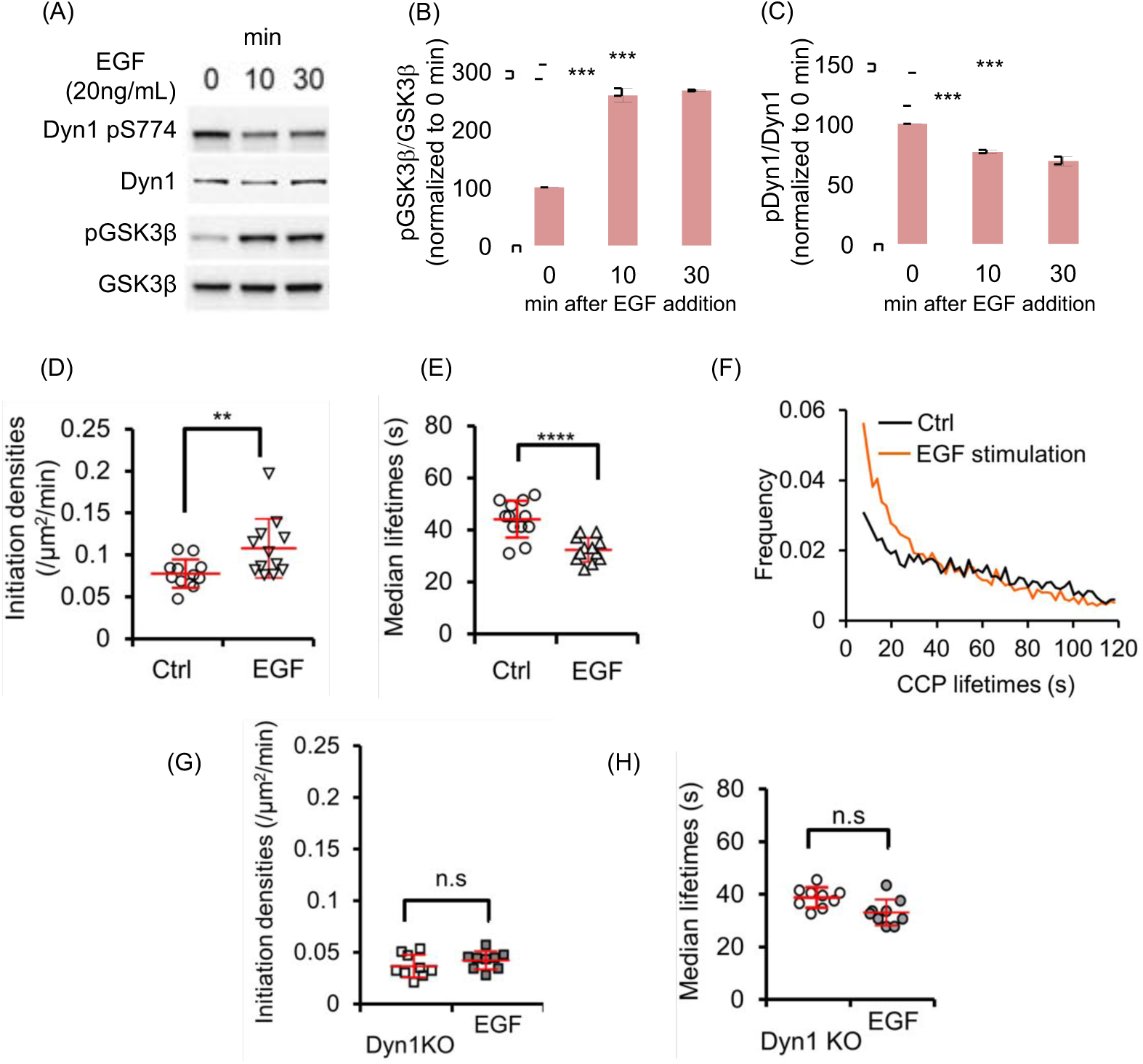
EGF stimulation alters CCP dynamics in a Dyn1-dependent manner. (A) Immunoblot analysis of changes in phosphorylation state of Dyn1 S774 and GSK3β upon EGF stimulation of parental A549 cells. (B,C) Quantification of the EGF-triggered changes in phosphorylation state (i.e. ratio of phosphorylated/ total protein) of GSK3β and Dyn1 (mean ± std. dev. of n=3 experiments, data are normalized to 0 min time point). CCP initiation densities (D), median lifetimes (E) and the lifetime distribution (F) of bona fide CCPs in serum starved A549 cells before (Control, Ctrl) or after incubation with EGF (20 ng/ml) for 10 min prior to imaging. CCP initiation densities (G) and median lifetimes (H) in serum-starved Dyn1^KO^ A549 cells before (Ctrl) or 10 min after incubation with EGF (EGF).

## Discussion

Our experiments provide further evidence that Dyn1, in addition to its well-studied roles in membrane fission during synaptic vesicle recycling, also has non-canonical functions as a regulator of the earliest stages of CME in non-neuronal cells. As in neurons, Dyn1 activity is negatively regulated by constitutive phosphorylation, and activated by dephosphorylation. When studied at endogenous levels of expression, we show that Dyn1 and Dyn2 have distinct functions in CME, reflected in their quantitatively and qualitatively different recruitment to CCPs. While Dyn1 is expressed at very high levels in the brain, it is, like Dyn2, also widely expressed albeit at lower levels in all tissues and cells [2]. Importantly, we show that acute activation of even low, nearly undetectable, levels of Dyn1 can increase the rates of CCP initiation and maturation to accelerate CME. These effects of Dyn1 activation are not accompanied by a burst of recruitment prior to CCV formation, and thus are likely mediated by unassembled Dyn1 tetramers associated with CCPs.

As in neurons [33], Dyn1 is constitutively inactivated in non-neuronal cells by phosphorylation at S774 in the PRD by GSK3. Acute chemical inhibition of GSK3β activates Dyn1 to alter CCP dynamics and increase the rate of CME. While GSK3β has numerous substrates, we show that Dyn1 is both necessary and sufficient to account for the effects of GSK3β inhibition on CCP dynamics and CME. Specifically, the effects of GSK3β inhibition on CME are dependent on Dyn1, but not Dyn2 expression, and Dyn1^KO^ cells reconstituted with a nonphosphorylatable mutant of Dyn1 show increased rates of CCP initiation and maturation that phenocopy the effects of GSK3β inhibition.

When activated, either by GSK3β inhibition or by mutation of S774 and S778 to alanine, Dyn1exhibits burst recruitment at late stages of CME and partially substitutes for Dyn2 activity. Yet, CME remains dependent on Dyn2, indicating that the two isoforms play functionally distinct roles in CME.

A direct comparison of the *in vitro* properties of Dyn1 and Dyn2 established that they differ in their curvature generating/sensing properties [14]. While Dyn1 is an efficient curvature generator that is able to tubulate and catalyze fission from planar lipid templates, Dyn2 is a curvature sensor that is able to catalyze membrane fission of highly curved lipid templates, but requires the synergistic activity of curvature-generating N-BAR domain-containing accessory factors to drive curvature generation and fission from planar templates [14, 15]. Strikingly, these biochemical differences could be ascribed to a single residue (Y600 in Dyn1, L600 in Dyn2) encoded within hydrophobic loops of the curvature-generating PH domain of dynamin [14]. Based on these biochemical differences, it was suggested that the unique properties of Dyn2 might enable this isoform to monitor CCP maturation and to catalyze fission only after the development of a narrow membrane neck connecting deeply invaginated CCPs to the plasma membrane. Unexpectedly, the findings presented here and elsewhere [23, 35, 39], establish that Dyn1 uniquely functions to regulate the earliest stages of CME, including the rate of CCP initiation and maturation.

Activation of Dyn1 also altered the shape of the lifetime distribution curve for CCPs from a broad Rayleigh-like distribution with a distinct peak at ∼30s to a more exponential distribution. We have previously suggested that the Rayleigh-like shape reflects, rate-limiting regulatory processes operating during the first 30s of CCP progression [20, 22, 38]. It is possible that due to its curvature-generating ability, and/or through interactions with other partner proteins, Dyn1 activation accelerates these complex early processes of CCP maturation.

Although Dyn1 and Dyn2 exhibit >80% sequence identity within their GTPase, middle, PH domains and GED, previous studies of the cellular activities Dyn1/Dyn2 chimeras have nonetheless revealed striking isoform-specific functional differences conferred by both the PH and GTPase domains [48, 49]. Most divergent among mammalian dynamin isoforms is the PRD, which functions to mediate interactions with numerous SH3 domain-containing binding partners, and has been shown to target dynamin to CCPs [40]. Earlier comparative studies of Dyn1 and Dyn2 [13], as well as Dyn1/Dyn2 PRD chimeras expressed at near endogenous levels [14], have shown that Dyn2 is more efficiently recruited to CCPs in a PRD-dependent manner. However, these studies did not take into account the negative regulation of Dyn1 by GSK3β phosphorylation. Here we reproduce and extend these findings by showing that the differential recruitment of Dyn1 is not due to phosphorylation of its PRD, at least on S774 or 778. Indeed, the recruitment of Dyn1 to CCPs was not significantly enhanced by GSK3β phosphorylation or when S774/S778 were mutated to nonphosphorylatable alanines. Thus, surprisingly, the effects of activated Dyn1 on CCP dynamics appear to occur independent of detectably enhanced recruitment to CCPs. It will be important to identify isoform-specific binding partners for Dyn1 and Dyn2 in non-neuronal cells.

To date, most dynamin binding partners, including endophilin, amphiphysin, and intersectin have been identified in brain lysates in which Dyn1 is highly expressed and may play a specialized function in rapid synaptic vesicle recycling. Thus, it is perhaps not surprising that siRNA knockdown of these dynamin binding partners in non-neuronal cells has only mild effects on, primarily, Dyn2-dependent TfnR endocytosis. Further studies are needed to identify essential non-neuronal effectors of both Dyn2 and Dyn1 function in CME.

Unexpectedly, our data suggests that SNX9, which was first identified as a major binding partner of Dyn2 in HeLa cells [45], interacts most strongly with dephosphorylated Dyn1, and that the effects of Dyn1 activation on early CCP maturation are dependent on SNX9. Published findings on SNX9 function in CME are enigmatic. Consistent with our findings in A549 NSCLC cells, siRNA-mediated knockdown of SNX9 has only mild effects on CME in several cell lines studied [46, 50]. While it has been suggested that these mild effects are due to redundant functions of the distantly related (40% sequence identity) SNX18 [51], this is not the case in all cell types [50, 51]. TIRFM studies on the recruitment of overexpressed SNX9-GFP to CCPs have also yielded differing results: it has been reported to be recruited coincident with [46], after [31] and before dynamin [51]. Interestingly, one study reported that SNX9 might linger at endocytic ‘hot-spots’ where it could function as an organizer of CCP nucleation [52]. Our results further suggest a more complex role for SNX9 at multiple stages of CME. SNX9 is not required for activated Dyn1-dependent increases in the rates of CCP initiation. However, it is required for the effects of Dyn1 activation on accelerating CCP maturation as indicated by the marked switch from Rayleigh-like to quasi-exponential CCP lifetime distributions, which is reversed by SNX9 knockdown. That SNX9 knockdown alone decreases the rate of CCP initiation and slows CCP maturation in cells expressing Dyn1^WT^, suggests other, potentially Dyn2-dependent and/or independent functions in CME. More work is required to define both the multiple functions of SNX9 in CME and to identify Dyn1-specific binding partners required for CCP initiation.Recent studies have shown that Dyn1 is upregulated and/or activated in several cancer cell lines [35, 39], leading to the suggestion that Dyn1 might function as a nexus between signaling and CME [2]. Here we show that Dyn1 can be activated downstream of EGFR to alter CCP dynamics. Previous studies showed that TRAIL-activated death receptors can activate Dyn1 to drive their selective uptake via CME [35]. Similarly, elegant studies on clathrin-mediated endocytosis of the G-protein coupled ‐adrenergic receptors have shown that they alter the maturation kinetics of the CCPs in which they reside through delayed recruitment of Dyn2 [53]. These authors did not examine Dyn1 recruitment or function. Further studies will be needed to determine whether other signaling receptors can selectively alter the composition and/or maturation kinetics of CCPs in which they reside and, if so, whether these changes, as suggested by our present data, are at least in part Dyn1-dependent.

Based on our results and other recent findings [23, 35, 39] regarding Dyn1 function downstream of signaling in non-neuronal cells, it is surprising that Dyn1 knockout mice develop normally, can live for up to 2 weeks after birth, and exhibit primarily neuronal defects [12]. This could, in part be due to the redundant function of dynamin-3, as Dyn1/Dyn3-null mice exhibit a more severe phenotype and die within hours of birth [11]. However, we speculate that kinase-based activation of Dyn1 might function at the level of individual CCPs to foster their initiation and accelerate maturation, perhaps at a threshold of signaling not reached during normal development. Indeed, recent evidence has pointed to cargo-selective roles of Dyn1 in regulating CME and signaling in cancer cells [2, 35, 39]. Therefore, it might be interesting to probe Dyn1 or Dyn1/Dyn3 knockout mice for other, potentially more subtle, non-neuronal phenotypes related to signaling in health and disease.

## Materials and Methods

### Cell culture, vector preparation, transfection and culture perturbations

Non-small cell lung cancer cell lines A549 and H1299 were kindly provided by Dr. John Minna (The Hamon Center for Therapeutic Oncology, Depts. of Internal Medicine and Pharmacology, UTSW) and were grown in RPMI 1640 (Life Technologies) with 5% FBS at 37°C and 5% CO2 and imaged in a temperature controlled chamber mimicking similar culture conditions.

The retroviral expression vector pMIEG was a modified pMIB (CMV-IRES-BFP) vector encoding Dyn1 cDNA with N-terminal HA-tag and C-terminal eGFP fusion tag. Point mutations to introduce S774/8A in Dyn1 cDNA were performed by site-directed mutagenesis. The lentiviral expression vector pLVX-puro (Clontech) encoded clathrin light chain A (CLCa) N-terminally tagged with mRuby 2 [54] or SNAP-tag [55] spaced with a 6 amino acid GGSGGS linker. The constructs were assembled from PCR fragments of mRuby2, SNAP and CLCa (for primers see Table 1) in yeast as described below and subsequently cloned into pLVX-puro. Lentiviruses were generated in 293T cells following standard transfection protocols [56] and were used for subsequent infections. They were prepared to transduce fluorescently tagged (mRuby2 or SNAP tag) CLCa fused to its N-terminus. Infected cells were selected using 10μg/ml puromycin for 4 days, conditions under which uninfected cells perished. The cells were passaged for 2 weeks before imaging for CME analysis. Retroviruses were also generated in 293T cells and used to stably transduce eGFP tagged Dyn1^WT^, Dyn1^S774/8A^ and Dyn2^WT^ proteins. Gene transduction was performed by exposing A549 or H1299 cells to retrovirus-containing cell culture supernatants. After 2 rounds of infection transduction with viral containing media, the recipient cells were expanded and FACS sorted for eGFP levels comparable to endogenous Dyn1-eGFP in A549 cells.

Transfections for siRNA knockdown experiments were carried out using Lipofectamine 2000 or Lipofectamin RNAi-Max (Life Technologies), following manufacturer’s protocol. For siRNA mediated knockdown, approximately 2×10^5^ cells (H1299) or 3×10^5^ cells (A549) were plated in each well of a six well plate. 20 nmol siRNA was used per well and two rounds of transfection across 5 days was sufficient to achieve over 90% knockdown.

Perturbation of culture conditions by GSK3β inhibitor involved addition of 10 μM CHIR99021 (Sigma) to pre-warmed culture media and incubation of cells for 30 min before additional analysis. Growth factor stimulation was performed by adding 20 ng/ml EGF (Invitrogen) to pre-warmed, serum-free culture media. Cells were analyzed after 10 min of incubation with EGF.

### Generation of genome-edited cell lines

Genome edited A549 and H1299 cells were generated by editing Dyn1 and Dyn2 to carry fusion tags. For the endogenous labeling of Dyn1 and Dyn2 with fluorescent reporter proteins, we chose an approach based on site directed introduction of CRISPR/CAS9n targeted DNA breaks and template assisted homology driven repair. eGFP fused to Dyn1 at its C-terminus was generated by CRISPR/Cas9n nickase strategy targeting the end of exon 21 of the DNM1 gene, inserting the last 19 amino acids of splice isoform “a”, a seven amino acid linker [32], monomeric eGFP with a stop codon and the SV40 polyadenylation signal. In the donor plasmid, this inserted sequence was flanked by ∼950 base pair homology arms for homology driven repair (HDR). The + gRNA pair was designed using publicly available software [57] with oligos DNM1-Nuclease-A-f/ DNM1-Nuclease-A-r and DNM1-Nuclease-B-f/ DNM1-Nuclease-B-r, respectively (Supplemental Table 1). For assembly of the donor vector, the segments were amplified with oligonucleotides coding RP11-348G11 (BACPAC Resources Center, Children’s Hospital Oakland Research Institute, Oakland, CA) covering the end of human DNM1 gene was used as template for the left and right (http://crispr.mit.edu/) and prepared as described approximately 30 nucleotide overhangs. The bacterial artificial chromosome clone homology arms (**Fig 1A**). The left and right homology arms were amplified using primer pairs DNM1-LH-f/DNM1-LH-r and DNM1-RH-f/DNM1-RH-r, respectively (see Supplementary Table 1). The 19 C-terminal amino acids of splice isoform “a” and the linker sequence DPPVATL [32] were covered with oligonucleotides DNM1-C-assembly-f and DNM1-C-assembly-r and amplified with short primers DNM1-Cterm-f and DNM1-Cterm-r. The sequence coding for monomeric eGPF and the SV40 polyadenylation signal were amplified from plasmid peGFP-N1 (Clontech), which carried the A206K mutation [58] with primers DNM1-eGFP-f/DNM1-eGFP-r and DNM1-pA-f/DNM1-pA-r, respectively. The first and last primers (DNM1-LH-f, DNM1-RH-r) also included overhangs for the E coli/yeast shuttle vector pRS424 [59]. For the DNM2-mRuby2 donor vector, the homology arms were amplified from the published [34] DNM2-eGFP construct (gift from D. Drubin, University of California, Berkeley) with primers DNM2-LH-f/ DNM2-LH-r and DNM2-RH-f/DNM2-RH-r, respectively. The mRuby2 segment together with the linker sequence, DPPVATL [32]}, was amplified from pmRuby2-C1 (Addgene æ40260) [54]. The PCR products were purified on 1% agarose gels and extracted using standard protocols before transformation into YPH500 yeast cells [60]. Yeast transformation, plasmid extraction and plasmid validation were performed as described earlier [61]. The guide-RNA plasmids for DNM1-eGFP and donor vectors for both DNM1-eGFP and DNM2-mRuby2 are currently being made available from Addgene (Accession IDs not yet available).

For both the edits, the nCas9 nickase + gRNA pairs or the ZFN nuclease pairs were added at 1 μg DNA concentration and 2 μg of the donor plasmid was added to this mixture in OptiMEM (Life Technologies). This mixture was then added to 5.5 μl of Lipofectamine 2000 (Life Technologies) in 150 μl OptiMEM (Life Technologies), briefly vortexed and incubated at room temperature for 15 min. The mixture was then added to cells plated 12 h earlier at 70% confluency (∼3×10^6^ cells/well in 6 well dish) with freshly replaced media. Transfect containing media was replaced by pre-warmed fresh media and the cells were allowed to grow for the next 48 h and then passaged for expansion in a 10 cm dish. The expanded cells were sorted as eGFP (or mRuby2) gene-edited single cells into 96 well plates, 4 days after transfection using a FACSAria 2-SORP (BD Biosciences, San Jose, CA) instrument equipped with a 300-mW, 488-nm laser and a 100-μm nozzle. Clonal expansion ensued by incrementing the culture dish area and maintaining a minimum 50% cell confluency. Single clones were then assayed for edits by western blotting and cells positive for genome edits were expanded. In order to generate double genome edited A549 cells, The A549 clone, 2C8, with homozygous Dyn2-mRuby2 knock-in was chosen and subsequently edited for Dyn1 and cell selection was performed as before using 150 ul FACS preliminary screen followed by western blotting for validation.

Dyn1 KO H1299 cells were generated as previously described [23] and the same strategy was employed to generate A549 Dyn1 KO cells. Briefly, cells plated in 6 well were transfected with 1 μg each of sgRNAs and Cas9 nickase encoding plasmids and co-transfected with a 20^th^ of peGFP plasmid. eGFP positive cells were assumed to have harbored both the sgRNA guides and single cell sorted by FACS. In addition, Dyn2 KO cells were generated using a similar double nickase strategy with single-guide RNAs (sgRNAs) CGATCTGCGGCAGGTCCAGGTGG and CGCCGGCAAGAGCTCGGTGCTGG in the pX335 vector. Complete knockout of Dyn1 and 2 was validated by western blotting.

### Immunoprecipitation, pulldowns and subcellular fractionation

#### GST-SH3 pulldown

GST-Amph II SH3 pulldown involved lysing H1299 cells in lysis buffer (50mM Tris, 150mM NaCl, 1X Protease Inhibitor Cocktail (Roche)) containing 0.2% Triton X-100. Cells were dounced with 27.5 G syringe for about 20 times or until most of the cells were ruptured to release intact nuclei. The post-nuclear fraction (PNF) was obtained by spinning the lysate at 10,000 × g at 4**°**C for 10 min and collecting the supernatant. About 3 mg of PNF in 1ml volume was used for each pulldown. Addition of beads (∼20 μl) with bait protein (GST-Amph II SH3) in PNF followed by gentle rotation for 1 hr allowed binding of target proteins. The bound fraction was washed twice with lysis buffer containing 0.2% Triton-X-100 and the resulting beads were denatured using 2X Laemmli buffer (Bio-Rad), reduced with 5% β-mercaptethanol, boiled and run on SDS-PAGE gel of appropriate separation capacity (7.5% or 12%, based on the target protein size). The pulldowns were analyzed by western blotting.

#### Subcellular fractionation

Confluent A549 cells in a 60 mm dish were detached with 1 ml 10 mM EDTA at 37 °C for 10 min and washed with PBS by centrifugation, and then resuspended in 0.5 ml buffer 2 (25mM HEPES, 250mM sucrose, 1 mM MgCl_2_, 2mm EGTA, pH 7.4). The resuspended cells were lysed through 3 cycles of freeze-thaw (rapid freezing in liquid nitrogen and slow thawing in room temperature water). Cytoplasm and membrane portions were separated by 30 min ultracentrifugation at 110kg in a Beckman Coulter rotor(TLA55). Pellets were resuspended with 0.5 ml buffer 2 and both supernatant and pellets were solubilized in 0.5% Triton X-100 for 10 min on ice, and then precipitated with 10% TCA, followed by 2 rounds of 1 ml acetone wash. SDS-PAGE gel electrophoresis and Western blot were applied as described above [62].

#### GFP-nAb Immunoprecipitation

Confluent A549 cells in a 10 cm dish were detached with 2 ml 10 mM EDTA at 37 °C for 10 min and washed with PBS by centrifugation, and then resuspended and gently lysed for 15 min on ice with 2 ml buffer 3 (0.5% Triton X-100, 25mM HEPES, 150mM KCl, 1 mM MgCl_2_, 2 mM EGTA, 1X Protease Inhibitor Cocktail (Roche), 1X Phosphatase Inhibitor Cocktail, pH = 7.4). Lysates were centrifuged at 5000xg, 4 °C for 5 min to remove nuclei, and protein concentration in the post-nuclear fraction (PNF) was determined by Bradford assay. 0.5 mg of the PNF was added to 30 μl GFP-nAb agarose (Allele Biotech), rotated for 2 hours at 4 °C and then spun down at 2500xg, 4 °C, 2 min. The agarose was washed twice (1ml/each) with two different buffers to fulfill different experimental purposes: 1) to probe the dynamin interactors, the agarose was washed with buffer 3; 2) to probe dynamin self-assembly, salt concentration in buffer 3 was brought up to 300 mM to remove indirect dynamin-dynamin interactions. 10% cell lysate, which is used to determine immunoprecipitation efficiency, was precipitated with 10% TCA and washed twice with acetone from ‐20 °C freezer. The samples were applied to SDS page gel and Western blot for analysis.

### Total internal reflection fluorescence microscopy (TIRFM)

Cells expressing appropriate fluorophores were cultured overnight on an acid-etched and gelatin coated coverslip, placed in a well in 6 well plate. At the time of imaging, cells were checked for adherence and spreading. When imaging SNAP-tagged proteins, labeling was performed by incubating cells in 1 ml of fresh, pre-warmed media containing 1μl of pre-dissolved SNAP-CELL 647-SiR dye (NEB). After 30 min incubation under standard incubator conditions, the media was aspirated, washed twice with sterile PBS and re-incubated in fresh culture media. The coverslips were mounted on glass slides with spacers and sealed with the same media. In the event of adding growth factor or inhibitor, cells were pre-incubated for appropriate time and the coverslips were mounted as before with the treated media. The coverslips were then imaged using a 60x 1.49 NA Apo TIRF objective (Nikon) mounted on a Ti-Eclipse inverted microscope with Perfect Focus System (Nikon) equipped with an additional 1.8x tube lens, yielding at a total magnification of 108x. TIRFM illumination was achieved using a Diskovery Platform (Andor Technology). During imaging, cells were maintained at 37**°**C in RPMI supplemented with 5% fetal calf serum. Time-lapse image sequences were acquired at a penetration depth of 80 nm and a frame rate of 2Hz (three or two channels) or 1Hz (single channel) using a sCMOS camera with 6.5mm pixel size (pco.edge).

### Quantitative analysis of Imaging

The detection, tracking and analysis of all clathrin-labeled structures and thresholding to identify bona fide CCPs was done as previously described using the cmeAnalysis software package [22]. Briefly, diffraction-limited clathrin structures were detected using a Gaussian-based model method to approximate the point-spread function [22], and trajectories were determined from clathrin structure detections using the u-track software [37]. Subthreshold CLSs (sCLCs) were distinguished from bona fide CCPs as previously described, based on the quantitative and unbiased analysis of clathrin intensity progression in the early stages of structure formation [22, 63]. Both sCLSs and CCPs represent nucleation events, but only bona fide CCPs represent structures that undergo stabilization, maturation and in some cases scission to produce intracellular vesicles [22, 63]). We report the rate of bona fide CCP formation, distribution of CCP lifetimes as well as intensity cohorts as described previously [22]. We also report mean and maximum signal intensities in two or three channels for each individual CCP. These are average and maximum signal intensities for individual CCPs as they are extracted by the previously described analysis software [22]. The extraction of CCPs is achieved by a new function added to the cmeAnalysis software published in [22] that allows us to link the classification of events, CCPs or CLSs, to more sophisticated analysis intensity time courses and lifetime. In this study we focused merely on per-CCP mean and maximum intensity values, which were averaged per movie (1-5 cells) and finally presented as per-movie distributions covering 10-30 cells per experimental condition. Differences between conditions were assessed by comparison of the normal-distributed per-movie distributions using Student’s t-test and a threshold of p <0.01 to mark statistical significance.

### Statistical Analysis

Control and treatment datasets were statistically analyzed with two-tailed unpaired Student’s t-tests using Graphpad Prism 5.0 (Graphpad Software, La Jolla, CA), from which *p* values were derived (* *p* < 0.05, ** *p* < 0.01, *** *p* < 0.001, **** *p* < 0.0001). Error bars representing standard error of the mean (sem) for at least three independent experiments were calculated using Microsoft Excel.

### Receptor internalization (endocytosis) assay

An in-Cell ELISA approach was used to quantitate internalization of transferrin receptor (TfnR) and EGFR, as previously described [23], using either anti-TfnR mAb (HTR-D65) [64] or biotinylated-EGF as ligands. Cells were grown overnight in 96-well plates at a density of 2×10^5^ cells/well and incubated with 4 mg/ml of D65 or 20 ng/ml of biotinylated-EGF (Invitrogen) in assay buffer (PBS4+: PBS supplemented with 1 mM MgCl_2_, 1 mM CaCl_2_, 5 mM glucose and 0.2% bovine serum albumin) at 37°C for the indicated time points. Cells were then immediately cooled down (4°C) to arrest internalization. The remaining surface-bound D65 or biotinylated-EGF was removed from the cells by an acid wash step (0.2 M acetic acid, 0.2 M NaCl, pH 2.5). Cells were then washed with cold PBS and then fixed in 4% paraformaldehyde (PFA) (Electron Microscopy Sciences) in PBS for 30 min and subsequently permeabilized with 0.1% Triton X-100/PBS for 10 min. Internalized D65 was assessed using a goat anti-mouse HRP-conjugated antibody (Life Technologies), and internalized biotinylated-EGF was assessed by streptavidin-POD (Roche). The reaction was developed by a colorimetric approach with OPD (Sigma-Aldrich), and additional color development was stopped by addition of 50 μl of 5M of H_2_SO_4_. The absorbance was read at 490 nm (Biotek Synergy H1 Hybrid Reader). Internalized ligand was expressed as the percentage of the total surface-bound ligand at 4°C (i.e., without acid wash step), measured in parallel [23]. Well-to-well variability in cell number was accounted for by normalizing the reading at 490 nm with BCA readout at 560 nm.

## Acknowledgments

We thank members of the Schmid lab for helpful discussions and critically reading the manuscript, Phillipe Roudot for help with data analysis, Marcel Mettlen for help with microscopy, Carlos Reis generated the Dyn2^KO^ A549 cells and David Drubin generously provided the Zinc-finger nuclease constructs. These studies were supported by NIH R01 grants GM073165 to SLS and GD, GM42455 and MH61345 to SLS and Welch Foundation Grant I-1823 to SLS.

---

**Supplementary table 1:**
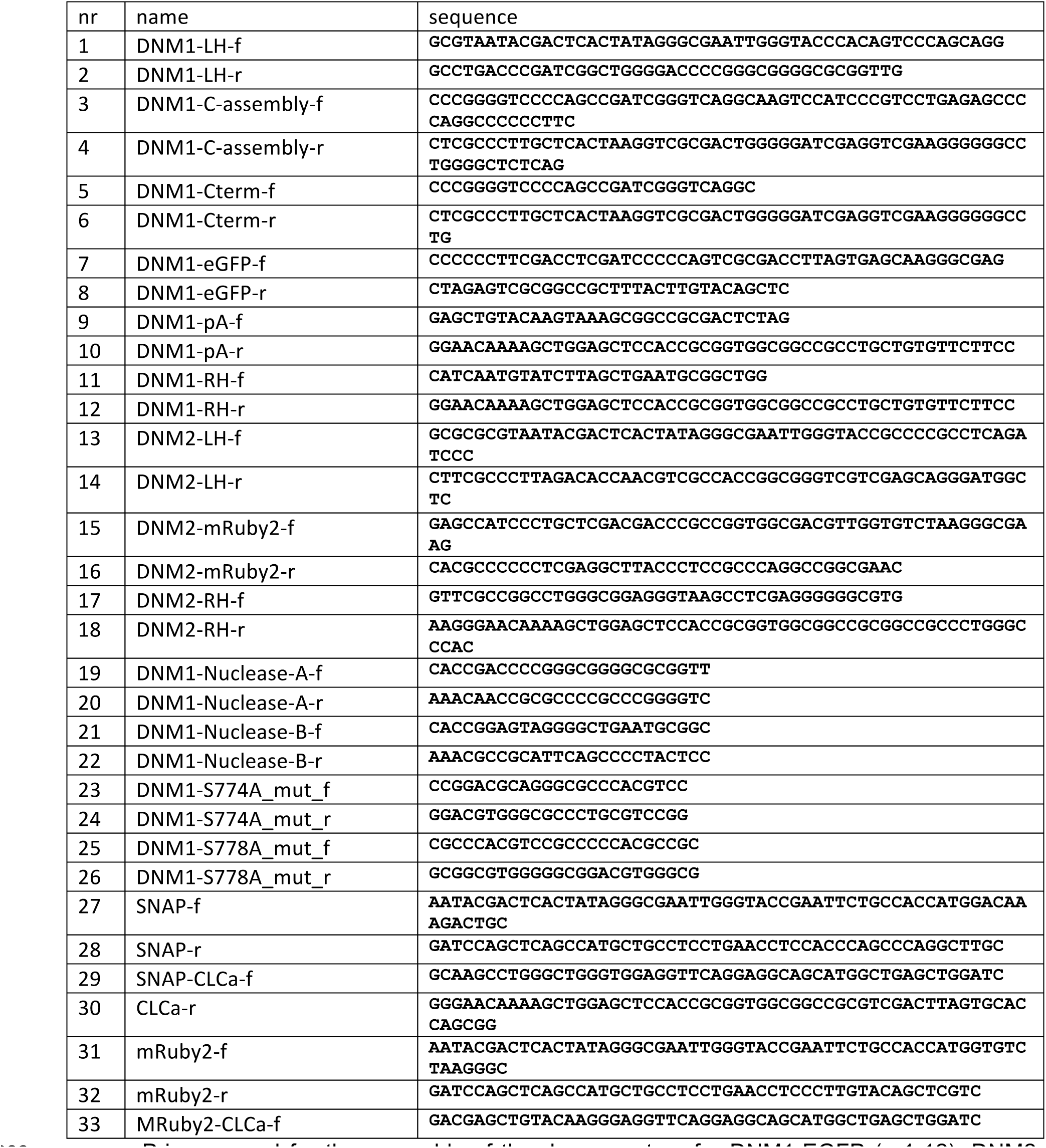
Oligonucleotides used for genome-editing, mutagenesis and fusion constructs Primers used for the assembly of the donor vectors for DNM1-EGFP (nr1-12), DNM2-mRuby2 (nr13-18), the nickase CRISPR/Cas9 guide RNA for DNM1 (nr19-22), DNM1 S774/8A mutagenesis (nr23 & 24) and for generation of the SNAP-CLCa and mRuby2-CLCa vectors (nr25-31). The donor DNA for DNM1-eGFP was assembled into the vector from five fragments: i) left homology (LH) arm, ii) DNM1 C-terminal 19 amino acids together with a seven amino acid linker (C-assembly), iii) monomeric eGFP (GFP), iv) SV40 polyadenylation signal (pA) and v) the right homology arm (RH), which were amplified with forward (f) and reverse (r) primers, as indicated.Likewise, the donor DNA for DNM2-mRuby2 was assembled into the vector from three fragments: i) the left homology arm (LH), ii) mRuby2 and iii) the right homology arm (RH). Primers 19-20 were used to generate the guides A and B for the nickase CRISPR-CAS9 guide RNA targeting DNM1. Primers 25-31 were used to generate CLCa-SNAP tagged lentiviral vectors.

**Figure S1.**
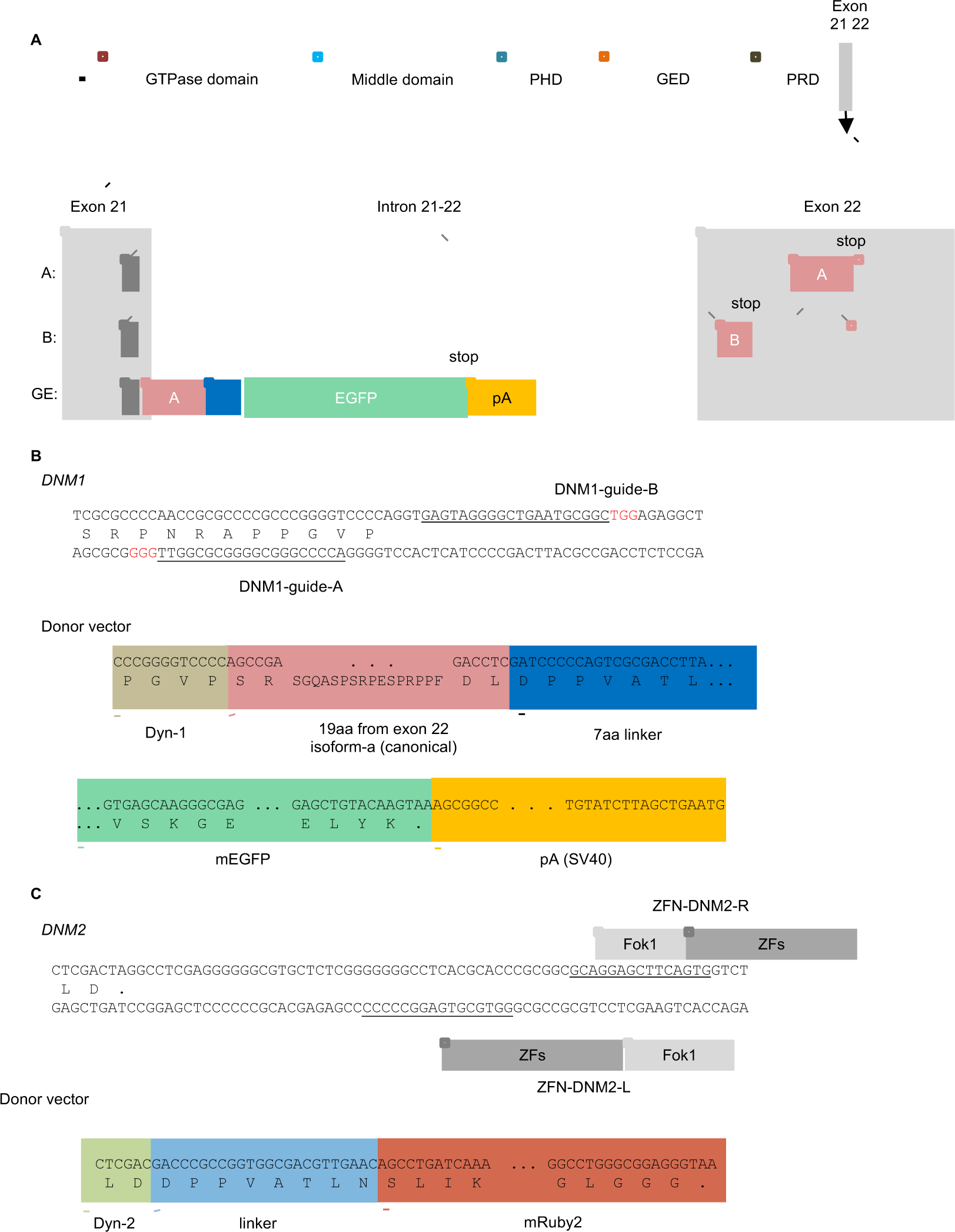
Design strategies for genome-edited H1299 and A549 cells. (A) Domain and genomic structure of DNM1 illustrating the C-terminal splice variants to illustrate splice variants A and B derived from exons 21 and 22 (filled red box) and 3′ UTR (open red box). To express the Dyn-1-EGFP fusion protein, the last 19 amino acids from splice variant A from exon 22 were introduced in frame in exon 21 (dark grey), followed by a 7 amino acid linker (blue), EGFP (green) and SV40 poly adenylation signal (orange). (B) Design of sgRNA guide A and B which targets the splice region in exon 21. The guide targeting sequences (underlined) and PAM sequences (red) are shown. For the donor vector, the DNA and amino acid sequences are shown for the junctions between exon 21, the inserted Dyn-1 C-term, the linker, EGFP and the poly adenylation signal. The color code is as in panel A. (C) Approach used for Zinc-finger nuclease-mediated genome-editing of DNM2, as previously described and the expected amino acid sequence for the Dyn2-mRuby2 fusion protein.

**Figure S2:**
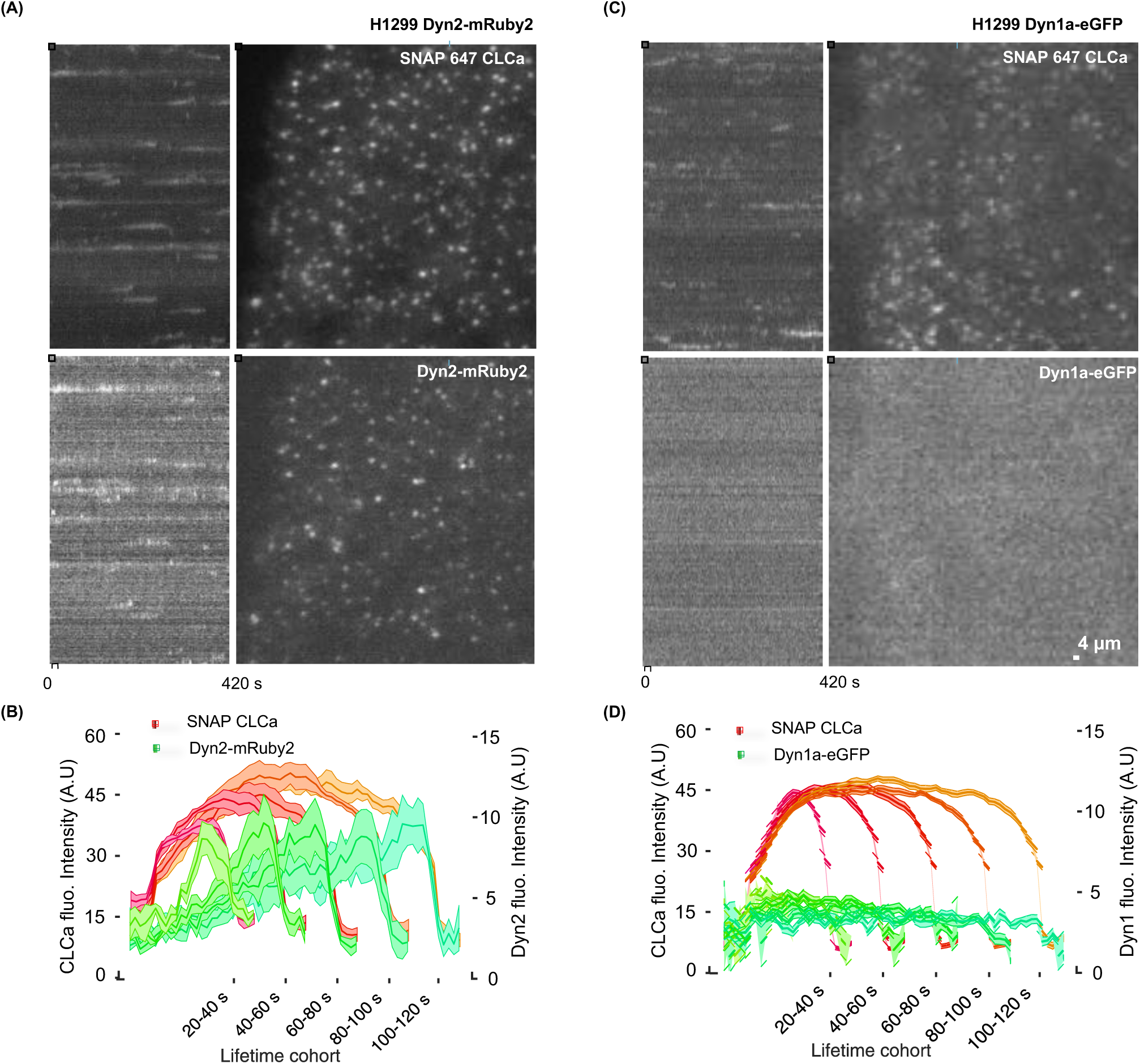
Isoform-specific differences in recruitment of Dyn1 and Dyn2 to CCPs. (A, C) Representative TIRF image and corresponding kymograph of dynamic SNAP(647)-CLCa-labeled clathrin coated pits and Dyn2-mRuby^end^ (A) or Dyn1a-eGFP^end^ (B) in genome-edited H1299 cells. (B, D) Corresponding quantification of the averaged intensities of CLCa and Dyn2-mRuby^end^ (B) or Dyn1a-eGFP^end^ (D) recruitment for the indicated lifetime cohorts. Data from 6647 CCPs from 5 independent movies, containing a total of 15 cells (B) and data from 74805 CCPs from 10 independent movies, containing a total of 29 cells (D).

**Figure S3:**
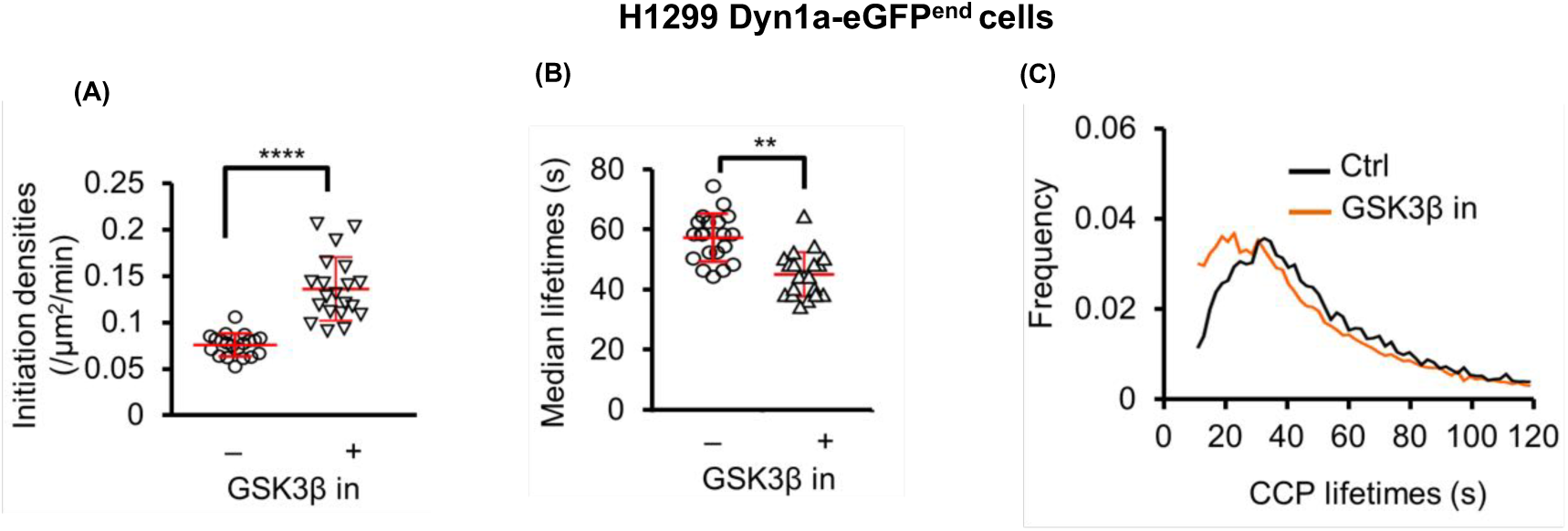
CCP dynamics in genome-edited Dyn1a-eGFP H1299 cells. (A) CCP initiation rates (B) CCP lifetimes and (C) lifetime distributions of all CCPs in H1299 cells genome edited to express endogenously-tagged Dyn1a-eGFP. Each point represents the value derived from a single movie, with 2-4 cells/movie. (** p≤0.01; ****p.0.0001)

**Figure S4:**
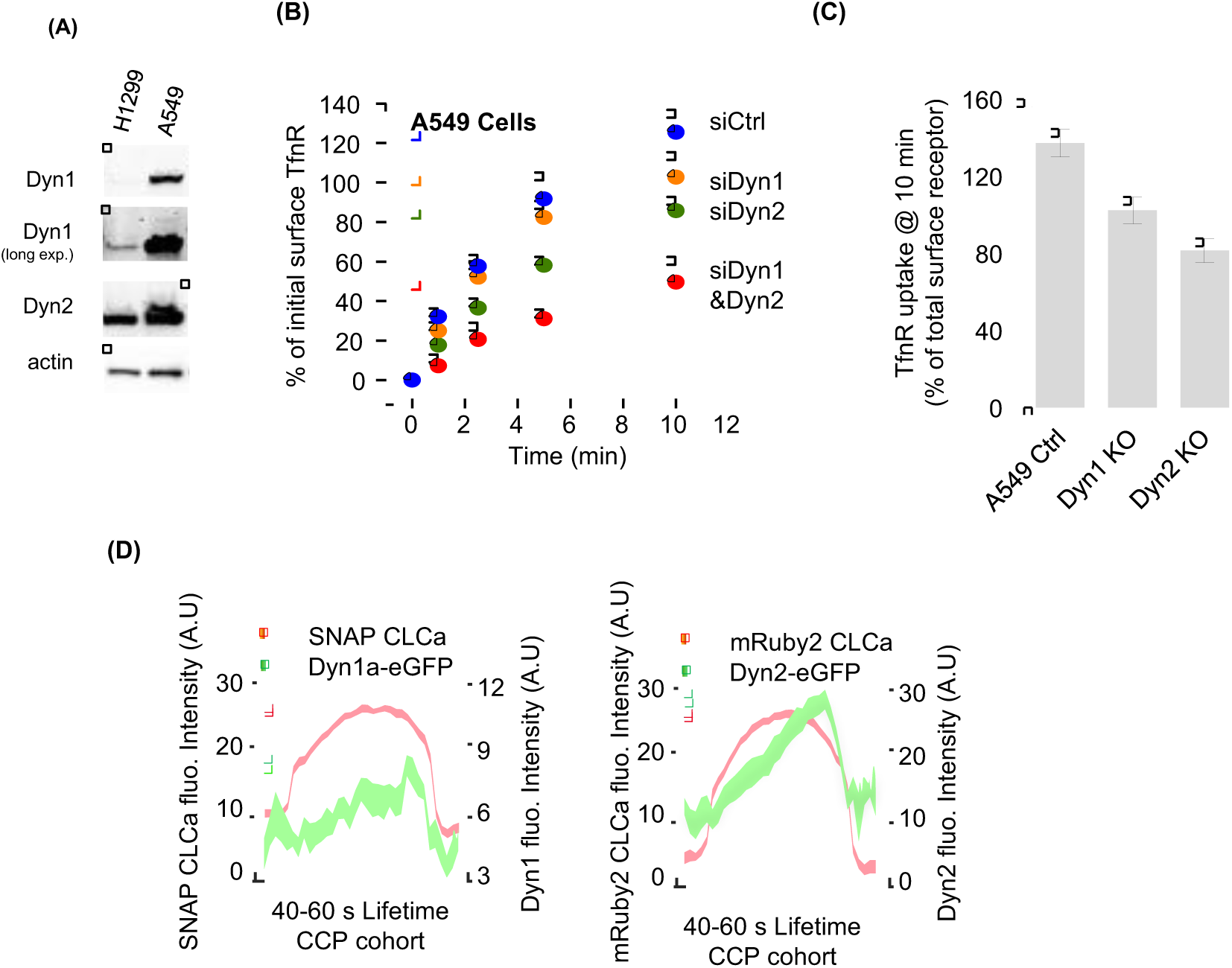
Characterization of TfnR endocytosis and dynamin-isoform recruitment in A549. (A) Differential expression of Dyn1 vs. Dyn2 in H1299 vs. A549 cells. In H1299 cells Dyn2 is expressed at ∼6-fold higher levels than Dyn1. In A549 cells Dyn1 is expressed at ∼5-fold higher levels Dyn2. (B) TfnR endocytosis in parental A549 cells treated with the indicated siRNAs. (C) TfnR uptake at 10 min in parental, Dyn1^KO^ and Dyn2^KO^ A549 cells. (D) Quantification of the average recruitment of Dyn1a-eGFP or Dyn2-eGFP to CCPs with lifetimes between 40-60s (4420 CCPs positive for Dyn1 and 3961 CCPs positive for Dyn2 were identified and analyzed from 11 movies containing 2-4 cells per movie), as in Figure 5E; however, the Dyn1a-eGFP data is rescaled to illustrate that Dyn1, like Dyn2 peaks at late stages of CME in these cells.

**Figure S5:**
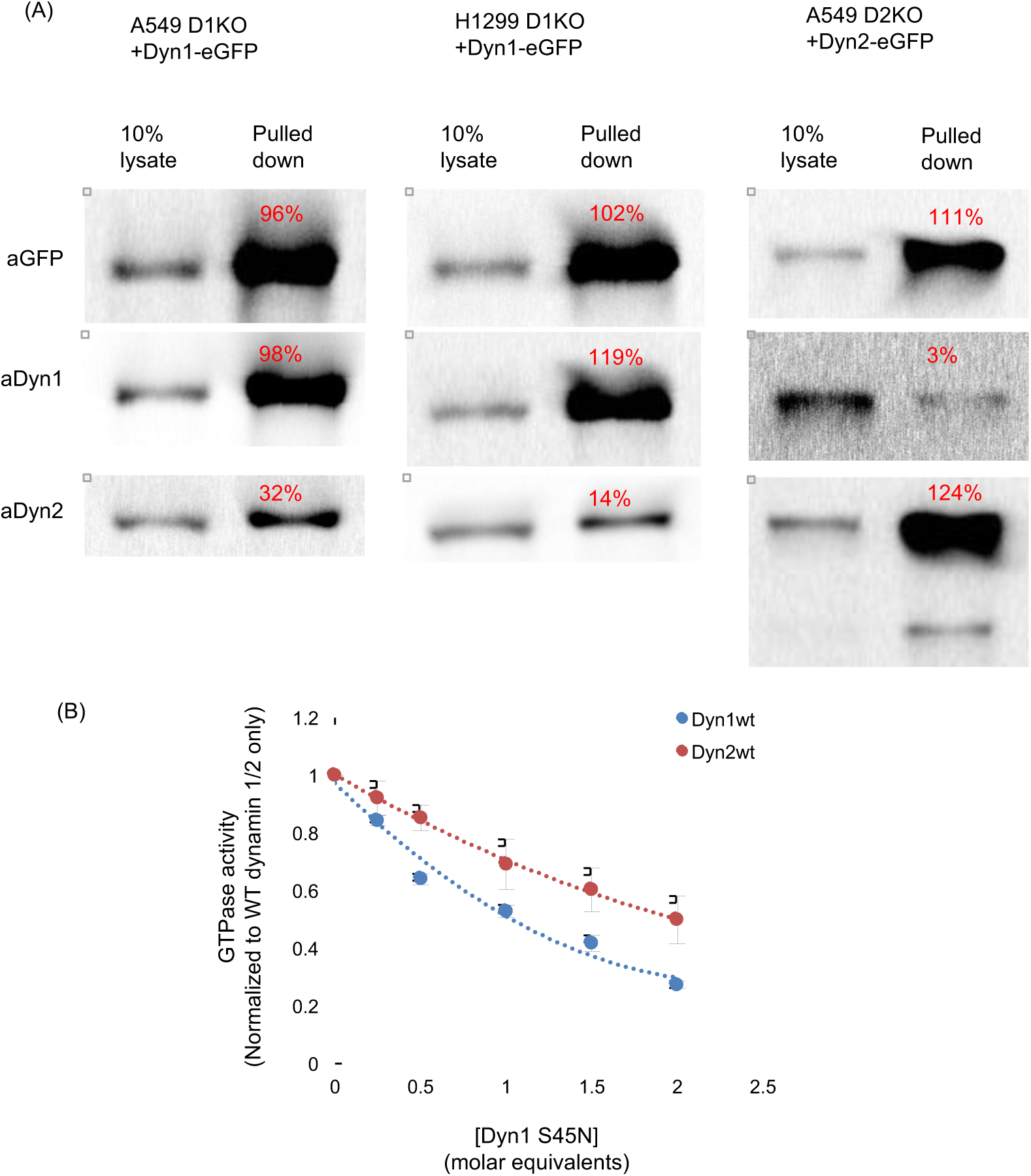
Dynamin isoforms only weakly co-assemble. (A) Western blots and quantification (red) of bands showing extent of pulldown of Dyn1-eGFP or Dyn2-eGFP using anti-eGFP nAb-beads and co-immunoprecipitation of the other isoform. Data are representative of 3 independent experiments. (B) The inhibition of assembly stimulated GTPase activity of Dyn1 (blue) or Dyn2 (red) in the presence of increasing concentrations of GTPase-defective Dyn1^S45N^, which will inhibit assembly-stimulated GTPase activity by co-assembling with WT-dynamin lipid nanotube templates.

## References

1. Mettlen M, Chen P-H, Srinivasan S, Danuser G, Schmid SL. Regulation of Clathrin-mediated Endocytosis. Ann Rev Biochem 2018;in press.

2. Schmid SL. Reciprocal regulation of signaling and endocytosis: Implications for the evolving cancer cell. J Cell Biol. 2017;216:2623–32.

3. Wideman JG, Leung KF, Field MC, Dacks JB. The cell biology of the endocytic system from an evolutionary perspective. Cold Spring Harb Perspect Biol. 2014;6:a016998.

4. Dergai M, Iershov A, Novokhatska O, Pankivskyi S, Rynditch A. Evolutionary Changes on the Way to Clathrin-Mediated Endocytosis in Animals. Genome Biology and Evolution. 2016;8:588–606.

5. Liu Y-W, Su AI, Schmid SL. The evolution of dynamin to regulate clathrin-mediated endocytosis Speculations on the evolutionarily late appearance of dynamin relative to clathrin-mediated endocytosis. Bioessays. 2012;34:643–7.

6. Schmid SL, Frolov VA. Dynamin: Functional design of a membrane fission catalyst. Ann Rev Cell Dev Biol. 2011;27:79–105.

7. Ferguson SM, De Camilli P. Dynamin, a membrane-remodelling GTPase. Nature Reviews Mol. Cell Biol. 2012;13:75–88.

8. Morlot S, Roux A. Mechanics of Dynamin-Mediated Membrane Fission. Ann Rev Biophysics. 2013;42:629–49.

9. Cao H, Garcia F, McNiven MA. Differential distribution of dynamin isoforms in mammalian cells. Mol. Biol. Cell. 1998;9:2595–609.

10. Antonny B, Burd C, De Camilli P, Chen E, Daumke O, Faelber K, et al. Membrane fission by dynamin: what we know and what we need to know. EMBO J. 2016;35:2270–84.

11. Raimondi A, Ferguson SM, Lou X, Armbruster M, Paradise S, Giovedi S, et al. Overlapping role of dynamin isoforms in synaptic vesicle endocytosis. Neuron. 2011;70:1100–14.

12. Ferguson SM, Brasnjo G, Hayashi M, Wolfel M, Collesi C, Giovedi S, et al. A selective activity-dependent requirement for dynamin 1 in synaptic vesicle endocytosis. Science. 2007;316:570–4.

13. Liu YW, Surka MC, Schroeter T, Lukiyanchuk V, Schmid SL. Isoform and splice-variant specific functions of dynamin-2 revealed by analysis of conditional knock-out cells. Mol Biol Cell. 2008;19:5347–59.

14. Liu Y-W, Neumann S, Ramachandran R, Ferguson SM, Pucadyil TJ, Schmid SL. Differential curvature sensing and generating activities of dynamin isoforms provide opportunities for tissue-specific regulation. Proc. Natl. Acad. Sci. (USA) 2011;108:E234–E42.

15. Neumann S, Schmid SL. Dual role of BAR domain-containing proteins in regulating vesicle release catalyzed by the GTPase, dynamin-2. J Biol Chem. 2013;288:25119–28.

16. Praefcke GJK, McMahon HT. The dynamin superfamily: Universal membrane tubulation and fission molecules? Nature Reviews Molecular Cell Biology. 2004;5:133–47.

17. Grassart A, Cheng AT, Hong SH, Zhang F, Zenzer N, Feng YM, et al. Actin and dynamin2 dynamics and interplay during clathrin-mediated endocytosis. J Cell Biol. 2014;205:721–35.

18. Hong SH, Cortesio CL, Drubin DG. Machine-Learning-Based Analysis in Genome-Edited Cells Reveals the Efficiency of Clathrin-Mediated Endocytosis. Cell Rep. 2015;12:2121–30.

19. Sever S, Muhlberg AB, Schmid SL. Impairment of dynamin’s GAP domain stimulates receptor-mediated endocytosis. Nature. 1999;398:481–6.

20. Loerke D, Mettlen M, Yarar D, Jaqaman K, Jaqaman H, Danuser G, et al. Cargo and dynamin regulate clathrin-coated pit maturation. PLoS Biol. 2009;7:e57.

21. Taylor MJ, Lampe M, Merrifield CJ. A feedback loop between dynamin and actin recruitment during clathrin-mediated endocytosis. PLoS Biol. 2012;10:e1001302.

22. Aguet F, Antonescu CN, Mettlen M, Schmid SL, Danuser G. Advances in analysis of low signal-to-noise images link dynamin and AP2 to the functions of an endocytic checkpoint. Dev Cell. 2013;26:279–91.

23. Reis CR, Chen PH, Srinivasan S, Aguet F, Mettlen M, Schmid SL. Crosstalk between Akt/GSK3beta signaling and dynamin-1 regulates clathrin-mediated endocytosis. EMBO J. 2015;34:2132–46.

24. Anggono V, Smillie KJ, Graham ME, Valova VA, Cousin MA, Robinson PJ. Syndapin I is the phosphorylation-regulated dynamin I partner in synaptic vesicle endocytosis. Nat Neurosci. 2006;9:752–60.

25. Huang Y, Chen-Hwang MC, Dolios G, Murakami N, Padovan JC, Wang R, et al. Mnb/Dyrk1A phosphorylation regulates the interaction of dynamin 1 with SH3 domain-containing proteins. Biochemistry. 2004;43:10173–85.

26. Damke H, Baba T, Warnock DE, Schmid SL. Induction of mutant dynamin specifically blocks endocytic coated vesicle formation. J Cell Biol. 1994;127:915–34.

27. Warnock DE, Baba T, Schmid SL. Ubiquitously expressed dynamin-II has a higher intrinsic GTPase activity and a greater propensity for self-assembly than neuronal dynamin-I. Mol Biol Cell. 1997;8:2553–62.

28. Merrifield CJ, Feldman ME, Wan L, Almers W. Imaging actin and dynamin recruitment during invagination of single clathrin-coated pits. Nat Cell Biol. 2002;4:691–8.

29. Ehrlich M, Boll W, Van Oijen A, Hariharan R, Chandran K, Nibert ML, et al. Endocytosis by random initiation and stabilization of clathrin-coated pits. Cell. 2004;118:591–605.

30. Rappoport JZ, Heyman KP, Kemal S, Simon SM. Dynamics of dynamin during clathrin mediated endocytosis in PC12 cells. PLoS One. 2008;3:e2416.

31. Taylor MJ, Perrais D, Merrifield CJ. A high precision survey of the molecular dynamics of mammalian clathrin-mediated endocytosis. PLoS Biol. 2011;9:e1000604.

32. Cocucci E, Gaudin R, Kirchhausen T. Dynamin recruitment and membrane scission at the neck of a clathrin-coated pit. Mol Biol Cell. 2014;25:3595–609.

33. Clayton EL, Sue N, Smillie KJ, O’Leary T, Bache N, Cheung G, et al. Dynamin I phosphorylation by GSK3β controls activity-dependent bulk endocytosis of synaptic vesicles. Nat Neurosci. 2010;13:845–51.

34. Doyon JB, Zeitler B, Cheng J, Cheng AT, Cherone JM, Santiago Y, et al. Rapid and efficient clathrin-mediated endocytosis revealed in genome-edited mammalian cells. Nat Cell Biol. 2011;13:331–7.

35. Reis CR, Chen PH, Bendris N, Schmid SL. TRAIL-death receptor endocytosis and apoptosis are selectively regulated by dynamin-1 activation. Proc Natl Acad Sci (USA). 2017;114:504–9.

36. Gaidarov I, Santini F, Warren RA, Keen JH. Spatial control of coated pit dynamics in living cells. Nature Cell Biol. 1999;1:1–7.

37. Jaqaman K, Loerke D, Mettlen M, Kuwata H, Grinstein S, Schmid SL, et al. Robust single-particle tracking in live-cell time-lapse sequences. Nat Methods. 2008;5:695–702.

38. Loerke D, Mettlen M, Schmid SL, Danuser G. Measuring the hierarchy of molecular events during clathrin-mediated endocytosis. Traffic. 2011;12:815–25.

39. Chen PH, Bendris N, Hsiao YJ, Reis CR, Mettlen M, Chen HY, et al. Crosstalk between CLCb/Dyn1-Mediated Adaptive Clathrin-Mediated Endocytosis and Epidermal Growth Factor Receptor Signaling Increases Metastasis. Dev Cell. 2017;40:278–88 e5.

40. Shpetner HS, Herskovits JS, Vallee RB. A binding site for SH3 domains targets dynamin to coated pits. J Biol Chem. 1996;271:13–6.

41. Ramachandran R, Surka M, Chappie JS, Fowler DM, Foss TR, Song BD, et al. The dynamin middle domain is critical for tetramerization and higher-order self-assembly. EMBO J. 2007;26:559–66.

42. Reubold TF, Faelber K, Plattner N, Posor Y, Ketel K, Curth U, et al. Crystal structure of the dynamin tetramer. Nature. 2015;525:404–8.

43. Liu YW, Mattila JP, Schmid SL. Dynamin-catalyzed membrane fission requires coordinated GTP hydrolysis. PLoS One. 2013;8:e55691.

44. Warnock DE, Hinshaw JE, Schmid SL. Dynamin self-assembly stimulates its GTPase activity. J Biol Chem. 1996;271:22310–4.

45. Lundmark R, Carlsson SR. Sorting nexin 9 participates in clathrin-mediated endocytosis through interactions with the core components. J Biol Chem. 2003;278:46772–81.

46. Soulet F, Yarar D, Leonard M, Schmid SL. SNX9 regulates dynamin assembly and is required for efficient clathrin-mediated endocytosis. Mol Biol Cell. 2005;16:2058–67.

47. Manning BD, Toker A. AKT/PKB Signaling: Navigating the Network. Cell. 2017;169:381–405.

48. Liu YW, Neumann S, Ramachandran R, Ferguson SM, Pucadyil TJ, Schmid SL. Differential curvature sensing and generating activities of dynamin isoforms provide opportunities for tissue-specific regulation. Proc Natl Acad Sci (USA). 2011;108:E234–42.

49. Soulet F, Schmid SL, Damke H. Domain requirements for an endocytosis-independent, isoform-specific function of dynamin-2. Exp Cell Res. 2006;312:3539–45.

50. Bendris N, Williams KC, Reis CR, Welf ES, Chen PH, Lemmers B, et al. SNX9 promotes metastasis by enhancing cancer cell invasion via differential regulation of RhoGTPases. Mol Biol Cell. 2016.

51. Posor Y, Eichhorn-Gruenig M, Puchkov D, Schoneberg J, Ullrich A, Lampe A, et al. Spatiotemporal control of endocytosis by phosphatidylinositol-3,4-bisphosphate. Nature. 2013;499:233–7.

52. Nunez D, Antonescu C, Mettlen M, Liu A, Schmid SL, Loerke D, et al. Hotspots organize clathrin-mediated endocytosis by efficient recruitment and retention of nucleating resources. Traffic. 2011;12:1868–78.

53. Puthenveedu MA, von Zastrow M. Cargo regulates clathrin-coated pit dynamics. Cell. 2006;127:113–24.

54. Lam AJ, St-Pierre F, Gong Y, Marshall JD, Cranfill PJ, Baird MA, et al. Improving FRET dynamic range with bright green and red fluorescent proteins. Nat Methods. 2012;9:1005–12.

55. Keppler A, Gendreizig S, Gronemeyer T, Pick H, Vogel H, Johnsson K. A general method for the covalent labeling of fusion proteins with small molecules in vivo. Nat Biotechnol. 2003;21:86–9.

56. Barde I, Salmon P, Trono D. Production and titration of lentiviral vectors. Curr Protoc Neurosci. 2010;Chapter 4:Unit 4 21.

57. Ran FA, Hsu PD, Wright J, Agarwala V, Scott DA, Zhang F. Genome engineering using the CRISPR-Cas9 system. Nat Protoc. 2013;8:2281–308.

58. Zacharias DA, Violin JD, Newton AC, Tsien RY. Partitioning of lipid-modified monomeric GFPs into membrane microdomains of live cells. Science. 2002;296:913–6.

59. Sikorski RS, Hieter P. A system of shuttle vectors and yeast host strains designed for efficient manipulation of DNA in Saccharomyces cerevisiae. Genetics. 1989;122:19–27.

60. Cermak T, Doyle EL, Christian M, Wang L, Zhang Y, Schmidt C, et al. Efficient design and assembly of custom TALEN and other TAL effector-based constructs for DNA targeting. Nucleic Acids Res. 2011;39:e82.

61. Gan Z, Ding L, Burckhardt CJ, Lowery J, Zaritsky A, Sitterley K, et al. Vimentin Intermediate Filaments Template Microtubule Networks to Enhance Persistence in Cell Polarity and Directed Migration. Cell Syst. 2016;3:252–63 e8.

62. van der Bliek AM, Redelmeier TE, Damke H, Tisdale EJ, Meyerowitz EM, Schmid SL. Mutations in human dynamin block an intermediate stage in coated vesicle formation. J Cell Biol. 1993;122:553–63.

63. Kadlecova Z, Spielman SJ, Loerke D, Mohanakrishnan A, Reed DK, Schmid SL. Regulation of clathrin-mediated endocytosis by hierarchical allosteric activation of AP2. J Cell Biol. 2017;216:167–79.

64. Schmid SL, Smythe E. Stage-specific assays for coated pit formation and coated vesicle budding in vitro. J Cell Biol. 1991;114:869–80.

